# Multi-kingdom microbial diversity and interaction landscapes in mosquitoes revealed by 5,163 individual meta-transcriptomes

**DOI:** 10.64898/2026.09.02.748197

**Authors:** Qin-Yu Gou, Wei-Chen Wu, Pei-Bo Shi, Geng-Yan Luo, Yuan-Fei Pan, Jing Wang, Yan Gao, Kai-Jie Liu, Hai-Long Zhao, Yun Feng, Kun Li, Wei-Hong Yang, De Wu, Shi-Jia Le, Gen-Yang Xin, Min-Wu Peng, Yu-Qi Liao, Chun-Hui Yang, Shi-Qiang Mei, Jia-Ming Huang, Jin-Xia Cheng, Xin Hou, Jian-Bin Kong, Xin-Xin Chen, Bing Zhang, Zi-Rui Ren, Jun-Hua Li, Xin Jin, Juan Wang, Tong-Qing An, Xin-Yi Huang, Jie Cui, John-Sebastian Eden, Gong Cheng, De-Yin Guo, Guo-Dong Liang, Edward C. Holmes, Zi-Qing Deng, Mang Shi, Da-Xi Wang

## Abstract

Mosquitoes are pathogen vectors embedded within diverse microbial ecosystems. However, the nature and interactions among their multi-kingdom microbiome remain poorly understood. We conducted a nationwide single-mosquito meta-transcriptomic survey of 5,163 mosquitoes representing 100 species across China, integrating viral discovery with marker-gene profiling of bacteria, archaea, fungi, and other eukaryotic microbes. From this, we identified 1,606 microbial species-level taxa, including extensive novel diversity, and revealed pronounced host species– specific organization of mosquito-associated communities. We detected 34 pathogens or potential pathogens of human or animal relevance, whose prevalence, abundance, host range, and geographic distribution defined distinct epidemiological patterns. Network analysis uncovered pervasive cross-kingdom microbial associations, including candidate antiviral relationships involving *Wolbachia* and other microbial taxa. Our study establishes a detailed view of the full-spectrum microbiome and provides a resource and conceptual framework for studying vector competence, pathogen emergence, and microbiome-informed mosquito-borne disease control.

## Introduction

Mosquitoes are among the most important vectors of human and animal pathogens, acting as ecological bridges between vertebrate hosts and a wide range of infectious agents. Species such as *Aedes aegypti* and *Aedes albopictus* drive the transmission of dengue^1^, Zika^2^, chikungunya^3^, and yellow fever viruses^4^, while *Culex* mosquitoes are major vectors of West Nile virus^5^, Japanese encephalitis virus^6^, and other arthropod-borne viruses (arboviruses). However, mosquitoes are not merely carriers of vertebrate pathogens as they harbor complex, multi-kingdom microbial communities that include non-vertebrate viruses, bacteria, archaea, fungi, and parasitic eukaryotes. Virological and metagenomic studies have also revealed an extensive diversity of insect-specific viruses (ISVs), highlighting that mosquitoes host rich viral ecosystems beyond classical arboviruses^7–13^. Beyond viruses, diverse bacterial symbionts (e.g., *Wolbachia*^14^, *Asaia*^15^, *Serratia*^16^), fungi (e.g., *Aspergillus*^17^, *Talaromyces*^18^), and parasitic eukaryotes (e.g., *Plasmodium*^19^, *Lambornella*^20^)—including nematodes^21^ and gregarines^22^—are widely present and exhibit either host-wide or tissue-specific (e.g., gut-associated) distributions. Hence, the mosquito-associated microbiome is inherently multi-kingdom in structure.

Despite our expanding knowledge of the mosquito microbiome, the complete assemblage of microbes has not been systematically resolved at the level of individual mosquitoes. Most existing studies rely on pooled samples that obscure natural variation and disrupt the linkage between host identity, co-infection patterns, and microbial load^8,23–27^. In addition, DNA-based approaches inherently fail to capture RNA viruses, limiting reconstruction of the complete microbial landscape^28–30^. Although recent single-mosquito meta-transcriptomic studies— including our own large-scale RNA sequencing efforts—have demonstrated the power of individual-level analysis, and several studies have shown that this approach can recover viral, prokaryotic, and eukaryotic signals within the same assay, systematic large-scale characterization of the mosquito full-spectrum microbiome remains limited^13,31,32^.

Of particular importance is that mosquito-associated microbes are not passive co-inhabitants, but instead form interaction networks that can shape vector competence. For example, *Wolbachia* suppresses multiple arboviruses and *Plasmodium*^14,33–36^, whereas bacteria such as *Serratia*^16^ and fungi such as *Talaromyces*^18^ can enhance viral susceptibility. Interactions also occur among non-viral microbes, including competitive exclusion between *Wolbachia* and *Asaia*^37^, and inhibition of *Plasmodium* transmission by Microsporidia MB^38^. These observations suggest that mosquito competence may not only be governed by individual pathogens, but by a network of interacting microorganisms. This underscores the need for a system-level understanding of the mosquito microbiome. In addition, these interaction networks can be leveraged for microbiome-based vector control strategies, including the exploitation of naturally occurring antiviral symbionts or the rational design of microbial consortia to suppress pathogen transmission in mosquito populations^14,34,39,40^.

Here, we applied large-scale single-mosquito meta-transcriptomics to define the mosquito full-spectrum microbiome across China. We simultaneously profiled RNA viruses, DNA viruses, bacteria, archaea, fungi, and other eukaryotic microbes, generating an integrated multi-kingdom map of mosquito-associated communities, and revealing cross-kingdom interaction networks linking viruses and other microbial groups. These data provide a comprehensive framework for understanding mosquito-associated microbial ecology and mosquito–microbe interactions, as well as their implications for pathogen transmission and control.

## Results

### Meta-transcriptomic Survey of Individual Mosquitoes in China

Between 2018 and 2025, we conducted a large-scale survey of mosquitoes across mainland China. Over 200,000 mosquito specimens were collected from 23 provinces, municipalities, and autonomous regions, encompassing diverse ecological and climate zones. To profile the full-spectrum microbiome of mosquitoes—including both pathogenic and symbiotic microorganisms such as viruses, bacteria, archaea, fungi, and other eukaryotic microbes—we selected 5,163 individual mosquitoes for meta-transcriptomic sequencing (Figure 1A). After quality control, the 5,163 libraries yielded a total of 1.423 trillion raw reads (mean: 275.6 million per sample). Following rRNA removal, 43.7 billion reads remained (mean: 8.5 million per sample), with 10.1 billion non-host reads remaining after the removal of host-derived reads (mean: 2.0 million per sample). Quality-controlled reads from each library were then *de novo* assembled into 140 million contigs, which were subsequently analyzed to characterize both the host and microbial components.

**Figure 1.**
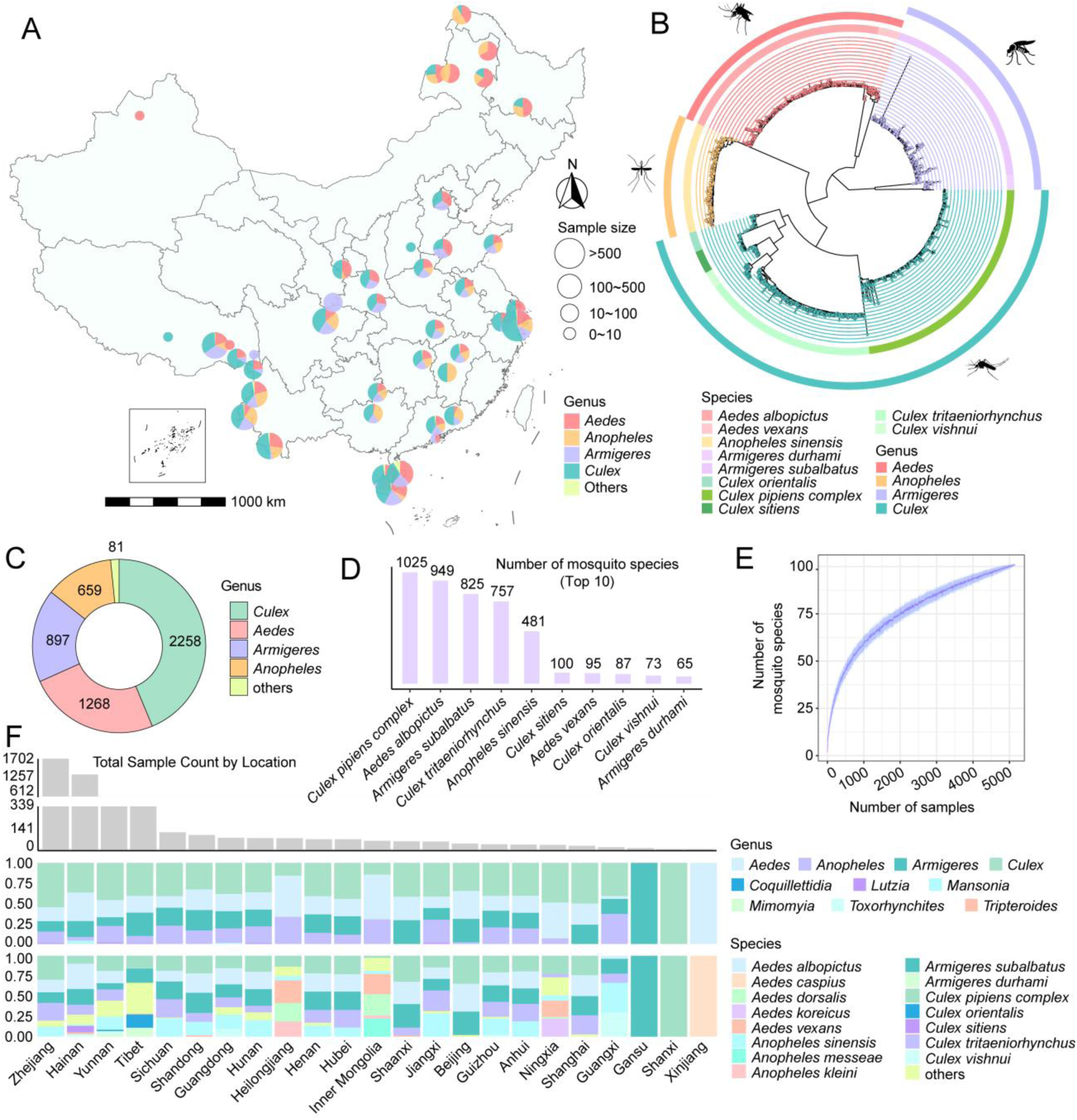
Sampling and phylogenetic profiling of 5,163 individual mosquitoes across China. **(A)** Sampling locations of 5,163 individual mosquitoes across China. Pie chart size indicates the number of samples per site, and colors represent mosquito genera. **(B)** Phylogenetic tree of the 10 most abundant mosquito species inferred from the mitochondrial *COX1* gene. The inner ring denotes species, and the outer ring denotes genera; distinct colors correspond to different taxa. **(C)** Relative composition of the dominant mosquito genera. Colors represent different genera. **(D)** Sample counts for the ten most abundant mosquito species. **(E)** Rarefaction curve showing mosquito species accumulation across samples. **(F)** Regional distribution of mosquito genera and species. Gray bars indicate total sample numbers per region; the middle and bottom panels show genus and species-level distributions, respectively, with colors representing distinct taxa.

Species identification based on the mitochondrial *COX1* gene revealed that these individuals represented 100 species across 10 genera, namely, *Aedes*, *Armigeres*, *Anopheles*, *Culex*, *Mansonia*, *Tripteroides*, *Toxorhynchites*, *Coquillettidia*, *Mimomyia*, and *Lutzia* (Figures 1B, 1C, and S1A; Table S1). The most prevalent mosquito species were *Culex pipiens complex* (n = 1,025), *Aedes albopictus* (n = 949), *Armigeres subalbatus* (n = 825), *Culex tritaeniorhynchus* (n = 757), and *Anopheles sinensis* (n = 481), together accounting for 78.2% of all samples (Figures 1D and S1B). Despite extensive sampling, the species discovery curve has not yet plateaued (Figure 1E). Geographic analysis further showed that these dominant genera and species were widely distributed nationwide, with most geographic regions harboring all four major genera and five dominant species, alongside region-specific taxa (Figure 1F).

### Revealing the Mosquito Microbiome

Contigs and reads were categorized into major biological groups, including the mosquito host, viruses, bacteria, archaea, fungi, and other eukaryotic microbes, and then systematically annotated and quantified to profile the full-spectrum microbiome. On average, the majority of non-rRNA reads originated from the mosquito host (93.47%). Among classified microbial groups, viruses accounted for the largest proportion of reads (62.59%), followed by bacteria (28.87%), fungi (4.60%), other eukaryotic microbes (3.83%), and archaea (0.12%) (Figure 2A). Across individual mosquitoes, viruses and bacteria dominated the microbial communities, with broadly consistent abundance patterns across species, except for *A. subalbatus* in which bacteria were substantially less abundant than viruses (Figure 2B).

**Figure 2.**
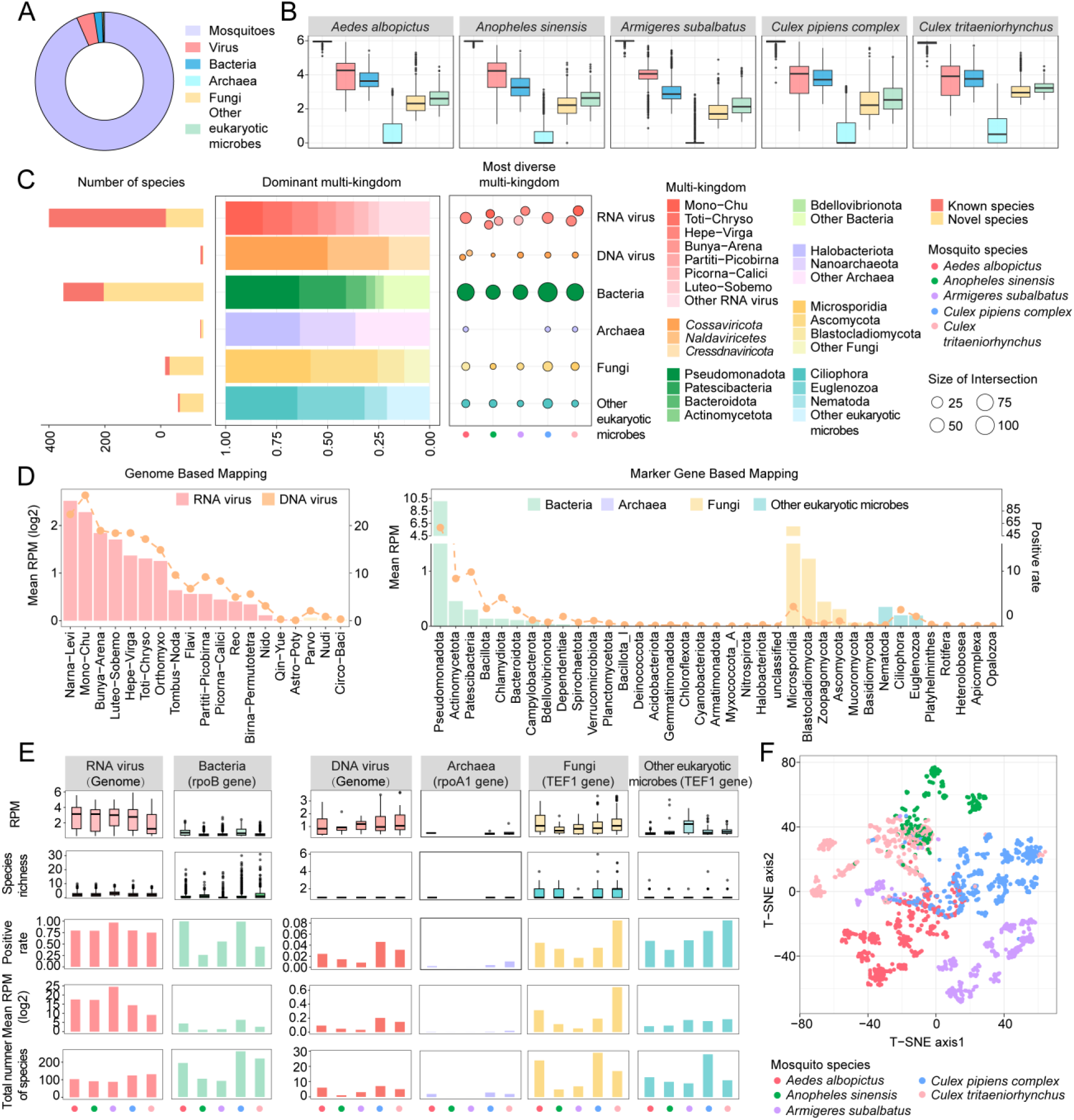
The multi-kingdom landscape of the mosquito microbiome. **(A)** Relative abundance of each organismal category within the mosquito microbiome: the mosquito host (purple), viruses (red), bacteria (blue), archaea (light blue), fungi (yellow), and other eukaryotic microbes (green). **(B)** Comparative abundance of each organismal category across the five dominant mosquito species. **(C)** Comparisons of diversity, measured as species richness, across different organismal categories. For each category, the left panel shows the total number of known (red) and novel (yellow) species; the middle panel presents relative richness across major viral supergroups or microbial phyla; and the right panel highlights the most diverse clades within each category across the five dominant mosquito species. **(D)** Comparisons of abundance and prevalence across organismal categories. For viruses, values were estimated based on genome or contig coverage; for prokaryotic and eukaryotic microbes, estimates were derived from marker genes. **(E)** Microbiome composition across the five dominant mosquito species. From top to bottom, panels depict total abundance distribution, richness distribution, prevalence, average abundance after logarithm transformation, and total number of species. **(F)** t-SNE ordination of individual mosquito microbiomes. Each point represents one mosquito sample, colored according to host species.

Using appropriate marker genes, we identified a total of 1,606 microbial species. After excluding non-mosquito-associated viruses, the remaining species comprised 553 mosquito-associated RNA viruses (134 novel), 10 mosquito-associated DNA viruses (2 novel), 501 bacteria (357 novel), 11 archaea (8 novel), 137 fungi (121 novel), and 91 other eukaryotic microbes (84 novel) (Tables S2, S3, and S4). We then assessed their diversity, abundance, and prevalence across all samples. Among mosquito-associated RNA viruses, the Mono-Chu, Toti-Chryso, and Hepe-Virga supergroups exhibited the greatest richness, whereas DNA viruses showed considerably lower diversity, with the *Cossaviricota* representing the most diverse clade (Figures 2C and S2). Consistent with richness patterns, the Mono-Chu group displayed the highest prevalence, whereas the Narna-Levi surpassed the Mono-Chu as the most abundant RNA supergroup, although most viruses within the Narna-Levi group were not mosquito-associated (Figure 2D). For cellular microbes, the phyla Pseudomonadota, Halobacteriota, and Microsporidia showed the highest richness, abundance, and prevalence among bacterial, archaeal, and fungal taxa, whereas the Nematoda had the highest abundance and Ciliophora the highest richness and prevalence among other eukaryotic microbial taxa, respectively, across all samples (Figures 2C and 2D).

We next compared microbiome profiles across the five dominant mosquito species at both broad and fine analytical levels. At the organismal category level, the overall structure of RNA viruses, assessed by total abundance, prevalence, and species richness, was largely conserved across species. In contrast, substantial interspecific variation was observed for fungi and DNA viruses: both their abundance and positivity rates were enriched in the *C. pipiens complex*, *C. tritaeniorhynchus*, and *A. albopictus*, but were markedly lower in *A. sinensis* and *A. subalbatus* (Figure 2E). To further examine whether these compositional differences translated into distinct microbiome architectures, we assessed microbiome similarity at the species-level OTU resolution using beta-diversity analyses. Accordingly, t-SNE ordination revealed clear mosquito species–specific clustering of microbiome composition (Figure 2F), a pattern that was further supported by PERMANOVA (*p* < 0.05).

### Composition and Diversity of Prokaryotic Communities in Mosquitoes

We characterized the mosquito-associated prokaryotic communities using *rpoB* (bacteria) and *rpoA1* (archaea) marker genes. Across all libraries, we identified 501 bacterial and 11 archaeal species-level OTUs (Figures 3A, S3, and S4A; Table S3), spanning 23 bacterial and 5 archaeal phyla. *Escherichia coli* was excluded due to its frequent occurrence as a sequencing contaminant. Overall, the prokaryotic community exhibited a highly uneven taxonomic distribution, with three bacterial phyla—Pseudomonadota (n = 180), Patescibacteria (n = 115), and Bacteroidota (n = 50)—accounting for 67% of all detected OTUs, and only a sparse representation of the remaining bacterial phyla and archaeal groups (Figures 3A and 3B). Strikingly, the majority of the taxa detected represented previously undescribed diversity: 71% of bacterial OTUs and 73% of archaeal OTUs represented putative novel species, and 33% of all prokaryotic OTUs shared <90% amino acid identity with known reference sequences (Figure 3B), underscoring a largely uncharacterized prokaryotic reservoir associated with mosquitoes. Next, we compared prokaryotic diversity across the ten mosquito species with the largest sample sizes. Prokaryotic richness varied substantially among mosquitoes, with *Culex orientalis*, *C. pipiens complex*, and *A. albopictus* exhibiting the highest richness, and *A. sinensis* and *A. subalbatus* harboring comparatively lower diversity (Figure 3C).

**Figure 3.**
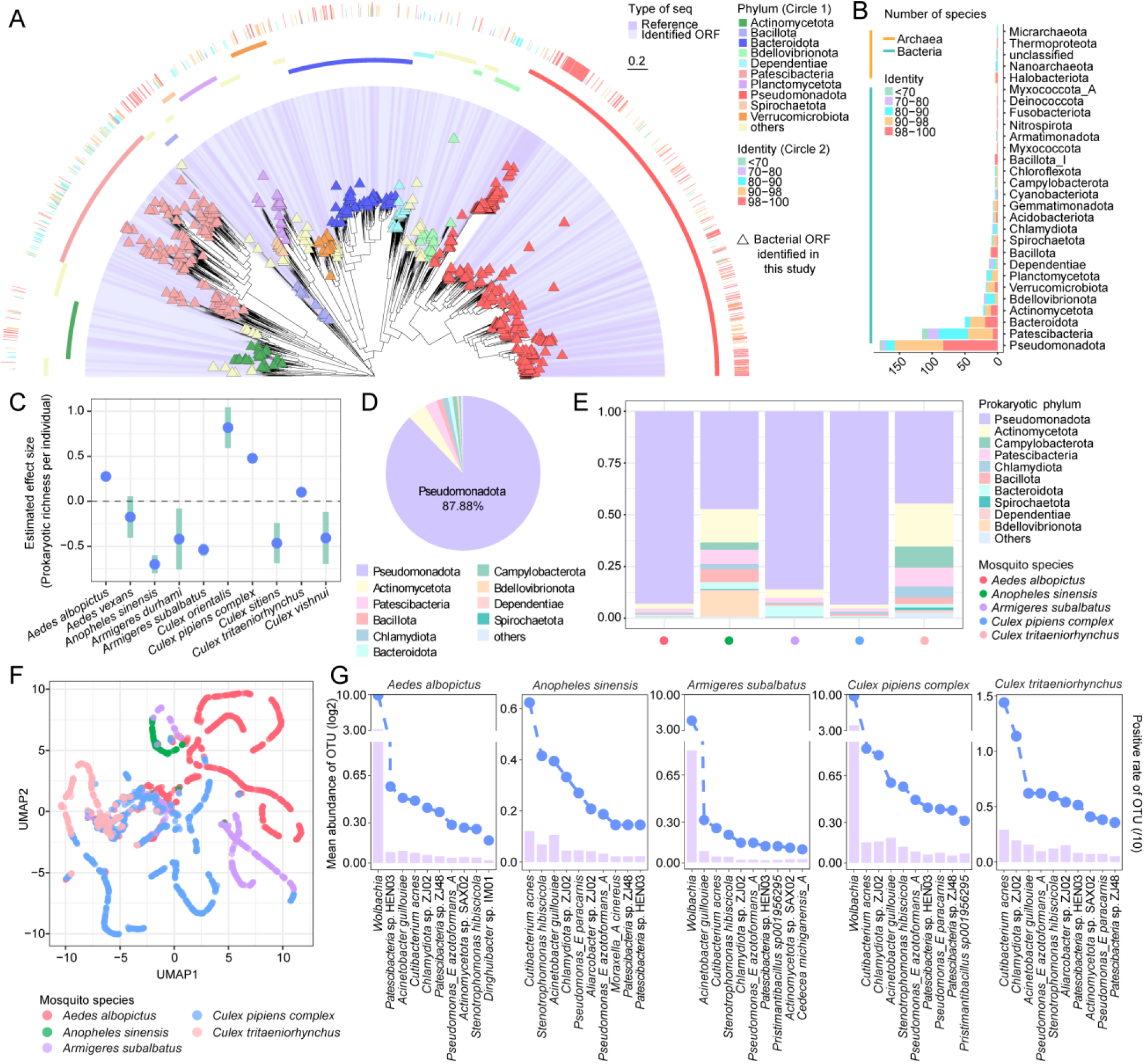
Diversity, composition, and host species-specific structure of mosquito-associated prokaryotic communities. **(A)** Phylogenetic tree of bacterial taxa estimated based on the *rpoB* protein. Triangles indicate *rpoB* sequences identified in this study, with colors representing bacterial phyla. Purple shading distinguishes sequence origin (reference versus newly identified). The inner annotation ring shows phylum-level taxonomic assignment, while the outer ring represents amino-acid sequence similarity, with colors corresponding to predefined similarity thresholds. **(B)** Distribution of species-level OTU richness across bacterial and archaeal phyla. Bar colors indicate amino-acid similarity levels. **(C)** Comparison of prokaryotic species richness among the top ten most abundant mosquito species. **(D)** Relative abundance of prokaryotic phyla across all mosquito samples. **(E)** Relative abundance of dominant prokaryotic phyla across the five most abundant mosquito species. Colors denote prokaryotic phylum, and colored circles at the bottom denote mosquito species. **(F)** UMAP ordination of dominant prokaryotic communities across the five most abundant mosquito species. Each point represents one mosquito sample, and colors indicate mosquito species. **(G)** Abundance (left axis, bars) and prevalence (right axis; dots and dashed line) of the top 10 most prevalent bacterial species-level taxa within the five most abundant mosquito species.

At the phylum level Pseudomonadota overwhelmingly dominated the mosquito-associated prokaryotic community, accounting for 87.88% of total abundance (Figure 3D). This was followed by Actinomycetota (3.97%), Patescibacteria (2.63%), Bacillota (1.19%), and other phyla (4.33%). A similar taxonomic composition was observed when the five dominant mosquito species were analyzed separately, although the relative proportions of minor phyla varied among host species, particularly beyond the two most dominant phyla, Pseudomonadota and Actinomycetota (Figure 3E). To validate marker-gene–based quantification, we compared *rpoB*-based abundance estimates against genome-wide coverage for *Wolbachia*, a dominant bacterium with deep sequencing representation (Figures S4B and S4C). The two approaches showed strong concordance (*r* = 0.9, *p* < 0.05; Figure S4D), supporting the robustness of marker-gene–based abundance estimation.

To characterize abundance at a finer taxonomic level, we further identified 61 dominant bacterial OTUs—defined as present in ≥20 libraries or >100 RPM—spanning eight phyla and collectively accounting for 93.48% of total bacterial reads (Figures S4E and S4F). Consistent with the overall microbiome patterns, UMAP ordination revealed clear host species–specific structuring of predominant prokaryotic communities (Figure 3F). Notably, dominant taxa varied markedly across mosquito hosts: *Wolbachia*-associated bacteria prevailed in *A. albopictus*, *A. subalbatus*, and *C. pipiens complex*, whereas *Cutibacterium acnes* was most prevalent in *A. sinensis* and *C. tritaeniorhynchus* (Figures 3G, S4F, and S4G). Despite these host-specific enrichments, the majority of dominant taxa (42/61) were shared across all five mosquito species (Figure S4F).

### Hidden Diversity of Eukaryotes in Mosquitoes

We next considered the eukaryotic component of the mosquito microbiome. Using *TEF1* protein-based taxonomic assignment and phylogenetic inference, we identified 137 fungal and 91 non-fungal eukaryotic species-level OTUs, spanning six and eight phyla, respectively (Figures 4A, 4B, and S5; Table S4). Among fungi, Microsporidia and Ascomycota exhibited the highest diversity, comprising 57 and 45 OTUs, respectively. In contrast, diversity among non-fungal eukaryotes was dominated by Ciliophora (32 OTUs) and Euglenozoa (30 OTUs) (Figures 4A, 4B, and S5).

**Figure 4.**
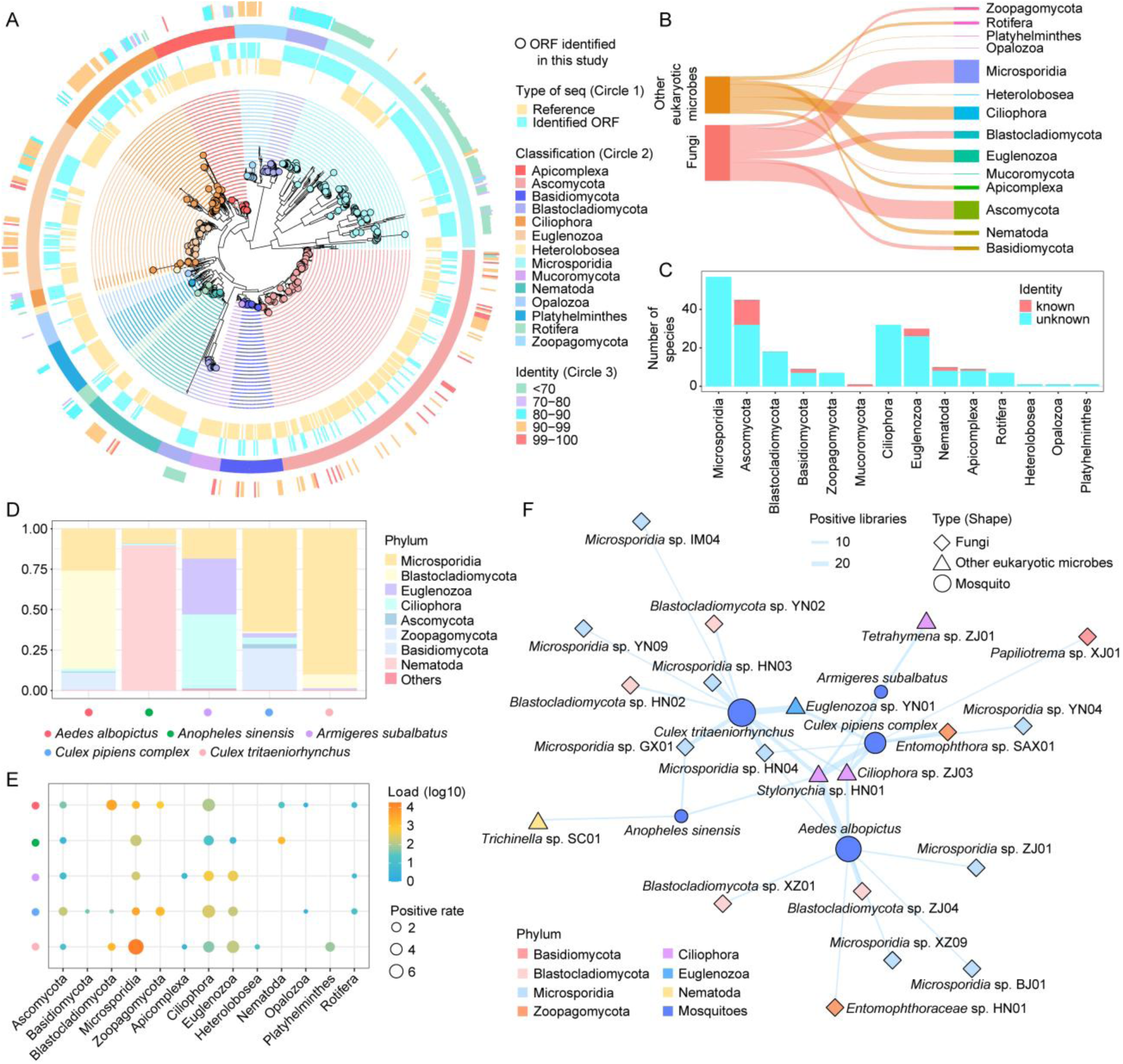
Diversity, host association, and cross-species distribution of fungal and other eukaryotic microbes within mosquitoes. **(A)** Phylogenetic tree of eukaryotic taxa based on the *TEF1* protein. Circles represent *TEF1* sequences identified in this study, with circle and shading colors representing eukaryotic phyla. The inner annotation ring indicates sequence origin (reference versus newly identified). The middle ring shows phylum-level taxonomic assignment, while the outer ring represents amino-acid sequence similarity, with colors corresponding to predefined similarity thresholds. **(B)** Species richness across eukaryotic phyla. The relative width of each shaded band reflects the number of species detected within each phylum. **(C)** Distribution of known and novel species across eukaryotic taxa. **(D)** Relative abundance of eukaryotic phyla across mosquito species. Colors denote eukaryotic phyla, and colored circles at the bottom indicate mosquito species. **(E)** Comparisons of abundance (colors) and prevalence (size) of eukaryotic phyla across mosquito species. **(F)** Cross-species distribution network of eukaryotic taxa. Circles represent mosquito host species, and other node shapes denote eukaryotic categories. Node colors correspond to distinct taxonomic classifications. Edge width indicates the number of positive libraries linking each microbial taxon to mosquito host species.

Consistent with the patterns observed for prokaryotic communities, the majority of detected eukaryotic taxa represented previously uncharacterized diversity. Specifically, 88.3% (121/137) of fungal OTUs and 92.3% (84/91) of non-fungal eukaryotic OTUs lacked close reference matches (Figure 4C), highlighting a largely unexplored eukaryotic reservoir associated with mosquito hosts. Comparison across the five dominant mosquito species revealed pronounced variation in eukaryotic abundance, with the highest load observed in *A. albopictus* and *C. tritaeniorhynchus*, and the lowest in *A. sinensis*. In contrast, species richness and alpha diversity did not reach statistical significance, although they tended to be higher in *C. tritaeniorhynchus* and *C. pipiens complex* and lower in *A. subalbatus* (Figure S6A).

At the phylum level, eukaryotic community composition differed markedly among mosquito hosts, with each species characterized by a distinct dominant phylum. Microsporidia predominated in the two *Culex* species (*C. tritaeniorhynchus* and *C. pipiens complex*), whereas Blastocladiomycota, Nematoda, and Ciliophora were dominant in *A. albopictus*, *A. sinensis*, and *A. subalbatus*, respectively (Figure 4D). These host-specific patterns were not only evident in relative abundance, but also in taxon prevalence across samples (Figure 4E). Notably, although Euglenozoa ranked only as the third most abundant lineage in *C. tritaeniorhynchus*, it exhibited a relatively high prevalence in this species (3.96%). Collectively, these pronounced host-associated patterns were further supported by t-SNE clustering, which revealed clear segregation of eukaryotic community profiles by mosquito species (PERMANOVA, *R²*= 0.08, *p* < 0.001; Figure S6B).

To compare composition and diversity at the species level, fungal and non-fungal OTUs were classified as dominant if detected in ≥20 libraries or exceeding 100 RPM. This yielded 30 dominant fungal OTUs spanning six phyla and 6 dominant non-fungal eukaryotic OTUs across three phyla. *Stylonychia* sp. HN01 was most prevalent in *A. albopictus* and *C. pipiens complex*, whereas *Microsporidia* sp. GX02, *Trypanosomatidae* sp. XZ01, and *Microsporidia* sp. HN03 were most dominant in *A. sinensis*, *A. subalbatus*, and *C. tritaeniorhynchus*, respectively (Figure S6C). Importantly, 16 dominant OTUs were shared across multiple mosquito species, indicating broad host ranges and suggesting the potential for cross-species transmission (Figure 4F).

### Mosquitoes Harbor an Epidemiologically Structured Spectrum of Pathogen-Related Lineages

We next delineated the subset of putatively pathogenic agents relevant to humans and other mammalian species. In total, 34 pathogens or potential pathogens were identified, including 6 arboviruses, 12 bacterial pathogens, 5 fungal pathogens, and 11 parasitic taxa (Figure S7). Among arboviruses, Tembusu virus and Getah virus showed >99% identity to known sequences, while additional diversity within *Rhabdoviridae* included both known and putative novel viruses related to agents implicated in human or animal disease. Based on the national notifiable infectious disease framework (National Catalogue of Human Infectious Pathogens), we identified 12 potential bacterial pathogens, including three taxa—most notably novel Bartonella lineages and known Rickettsia with the potential for arthropod-borne transmission, alongside clinically significant genera such as *Aeromonas*, *Bacillus*, *Providencia*, and *Vibrio*. We further detected five fungal pathogens (including *Alternaria alternata* and *Mucor circinelloides*) and 11 parasitic taxa spanning Nematoda, Apicomplexa, and Euglenozoa, including *Theileria orientalis*, *Toxoplasma*, multiple *Trypanosoma* and *Leishmania* lineages, and medically important nematodes such as *Brugia malayi*, *Loa loa*, and *Anisakis*-related species. This underscores the extensive and clinically relevant pathogenic sequence diversity within the mosquito microbiome.

To characterize the epidemiological structures of these pathogens, we quantified their abundance, prevalence, geographic distribution, and host breadth. Arboviruses were detected in 19 of 5,163 libraries, indicating a low prevalence, although they did reach high abundance within individual mosquitoes (up to 13,972.7 RPM) (Figure 5A). In contrast, bacterial pathogens—most notably *Providencia rettgeri*—were detected in the greatest number of libraries, spanned the widest geographic ranges, and may infect the broadest diversity of mosquito hosts (Figure 5A). Pathogen richness and composition also varied across mosquito species: *C. tritaeniorhynchus* showed the highest arboviral pathogen diversity, whereas *C. pipiens complex* contained many non-viral pathogens (Figure 5B). Overall, these two species harbored the richest pathogen repertoire. However, even sparsely sampled species, such as *Aedes caspius* (n = 10), exhibited substantial pathogen diversity (Figure 5C), suggesting that pathogen richness is not strictly driven by sampling depth.

**Figure 5.**
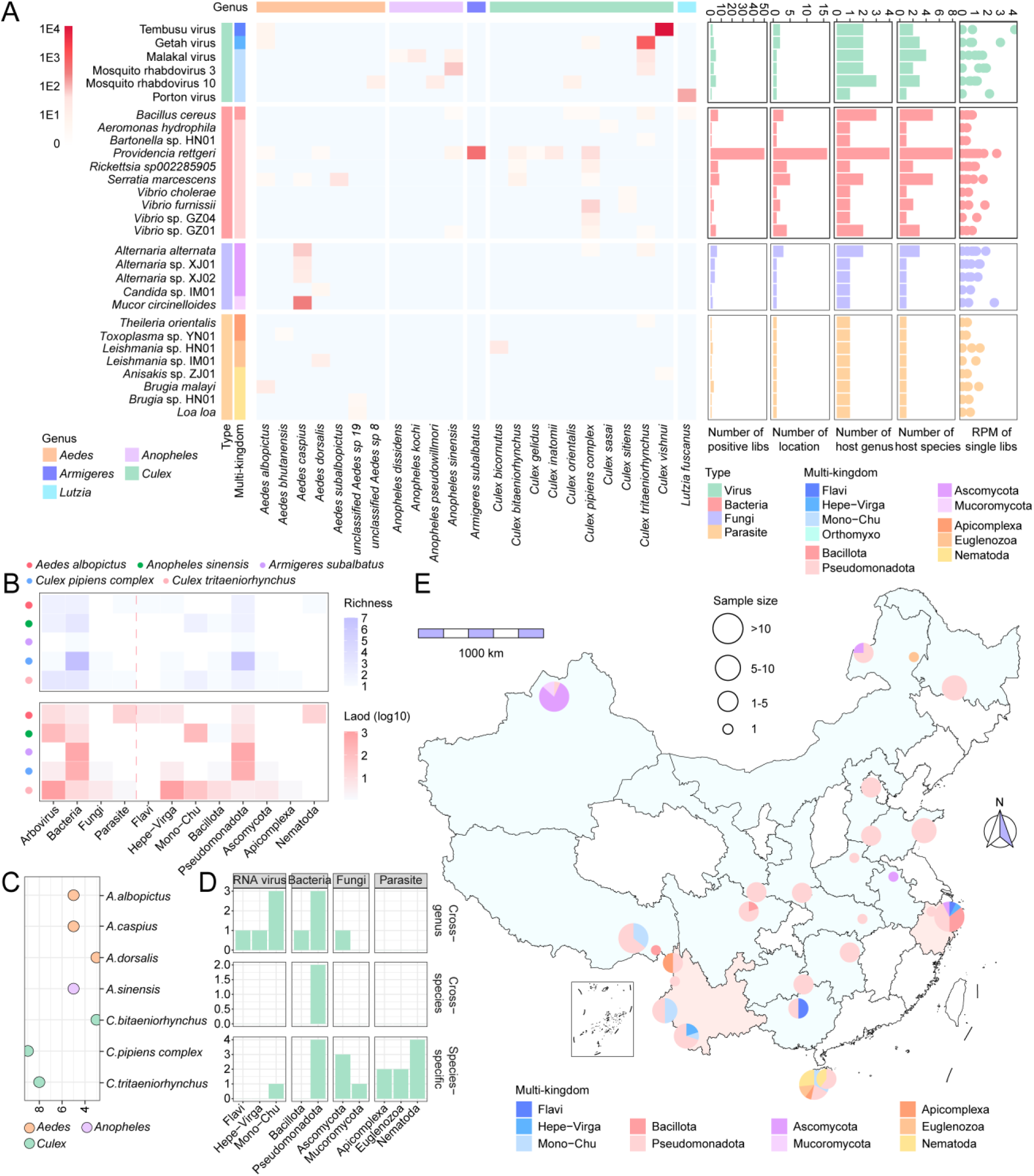
Pathogen landscape within mosquitoes. **(A)** Maximum load of pathogen-related taxa detected across mosquito species (heatmap), along with the number of positive libraries, sampling region, mosquito genera and species harboring each pathogen, and their loads per library (from the left to the right panel). Color intensity in the heatmap indicates pathogen load. Annotation colors of pathogens denote pathogen categories and taxonomic groups, while host annotations represent mosquito genera. Bar colors indicate pathogen categories. **(B)** Comparisons of pathogen richness and abundance across mosquito species. Colors represent pathogen richness in the upper panel and pathogen load in the lower panel. **(C)** Number of pathogen taxa detected in each mosquito species. **(D)** Summary of host-range patterns for pathogenic taxa, including cross-genus, cross-species, and species-specific occurrence. **(E)** Geographic distribution of pathogens across China. Pie size corresponds to the number of pathogen-positive samples, and colors indicate pathogen categories. Pale pink shaded regions indicate areas with relatively high pathogen diversity.

Cross-host occurrence was observed for 12 of 34 pathogens, including bacterial, viral, and fungal taxa (Figure 5D). For example, Getah virus was detected across multiple mosquito genera despite its low overall prevalence (0.06%), indicating broad host range. Geographically, southern China—particularly Yunnan, Hainan, and Zhejiang—harbored the highest pathogen diversity (Figure 5E), likely reflecting a combination of sampling intensity and regional ecological factors.

### Cross-Kingdom Interactions Structure the Full-Spectrum Mosquito Microbiome

To investigate interactions within the mosquito microbiome, we performed species-level Spearman correlation analysis based on abundance, identifying 148 significant positive and negative associations (Figures 6A and S8A). Associations were most frequent within bacteria (87 pairs), within RNA viruses (38 pairs), and between bacteria and RNA viruses (19 pairs) (Figure S8B). The majority of associations were positive (80.41%), with correlation strengths predominantly moderate (0.4–0.8), and stronger associations (|*r*| > 0.5) largely concentrated among bacteria–bacteria, RNA virus–RNA virus, and bacteria–RNA virus pairs (95.35%) (Figures 6B and 6C). These patterns indicate extensive coexistence and structured microbial co-occurrence within mosquitoes. Beyond broad trends, the network revealed pervasive cross-kingdom interactions spanning viruses, bacteria, fungi, and other eukaryotic microbes, including both positive and negative associations (Figure 6D). These findings suggest that the mosquito microbiome may operate as an ecological network rather than a collection of independent taxa.

**Figure 6.**
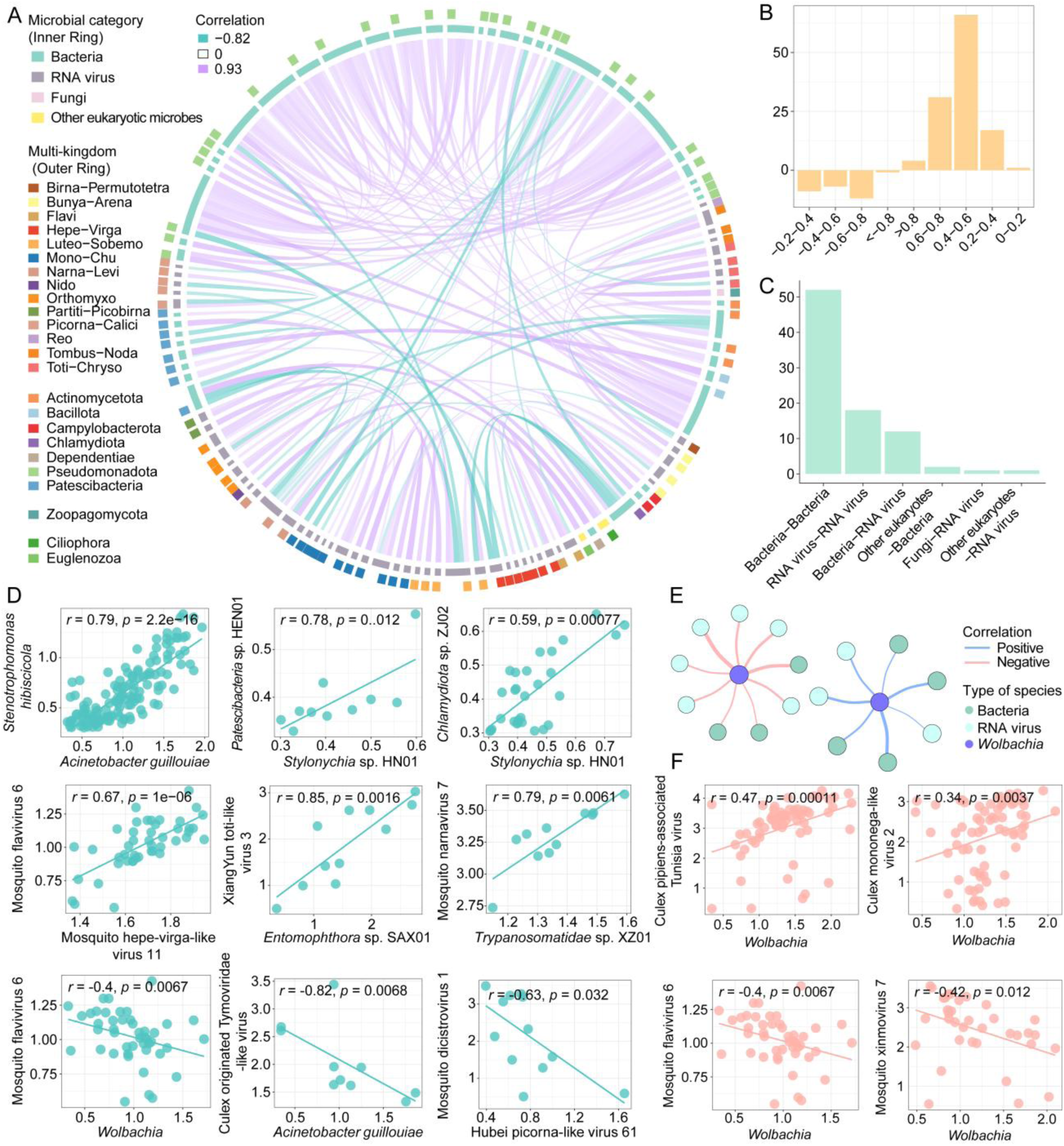
Associations within the mosquito full-spectrum microbiome. **(A)** Species-level correlation network. The inner annotation ring denotes microbial categories, and the outer ring indicates specific taxonomic groups (e.g., bacterial phyla and viral supergroups). Edge colors represent positive or negative correlations. **(B)** Distribution of Spearman correlation coefficient intervals. **(C)** Distribution of strong correlations (|*r*| > 0.5) among different microbial categories. **(D)** Pairwise correlations between species based on Spearman analysis. **(E)** *Wolbachia*-associated correlations (Spearman). **(F)** Correlations between *Wolbachia* and viral taxa.

Of most note we identified 16 potentially antiviral associations, including *Wolbachia* and *Acinetobacter guillouiae* that showed significant negative correlations with Mosquito flavivirus 6 and Culex originated Tymoviridae-like virus, respectively (Figure 6D). Given that *Wolbachia* is known to modulate arboviral infection, we further examined *Wolbachia*-associated relationships. We identified 16 significant *Wolbachia*-associated interactions, encompassing both positive (n = 3) and negative (n = 6) correlations with viral taxa (Figures 6E, 6F, and S8D). These context-dependent associations suggest a potentially complex role for *Wolbachia* within the broader microbial community. In addition, we identified four microbial taxa (including *Wolbachia* and *Opisthorchiidae* sp. YN01) that were negatively correlated with overall virome abundance, suggesting potential broad antiviral effects, although this requires further validation (Figure S8C). Collectively, these results reveal a densely connected, cross-kingdom interaction architecture within the mosquito microbiome and highlight candidate antagonistic and facilitative taxa that may influence viral abundance patterns and transmission dynamics.

## Discussion

Mosquitoes are major infectious disease vectors and are hosts to complex, multi-kingdom microbial communities, some of which may modulate or antagonize pathogen transmission. Here, through an expansive nationwide meta-transcriptomic survey of 5,163 individual mosquitoes, we define the mosquito full-spectrum microbiome and provide a multi-kingdom view of viruses, bacteria, archaea, fungi, and other eukaryotic microbes. This work substantially expands the known diversity of mosquito-associated microorganisms and establishes a foundation for studying microbial ecology and pathogen dynamics in vector systems. Importantly, the data generated represent one of the most comprehensive resources of mosquito-associated microbes, enabling future studies on host–microbe interactions, pathogen emergence, and microbiome-informed vector control.

A central finding of this study was the remarkable diversity and novelty of the mosquito microbiome. We identified over 1,600 microbial taxa, including 1,303 mosquito-associated species-level taxa, with more than 50% representing putative novel species, and an even higher proportion among cellular microbes (77%). Compared with previous DNA-based studies, which reported substantially fewer taxa, our analysis reveals a markedly expanded diversity landscape^41–45^. This is largely driven by meta-transcriptomic sequencing which, despite uneven genomic coverage, preferentially captures conserved and actively expressed genes, enabling reliable detection of low-abundance cellular organisms through well-covered marker loci^7,12,26,31,46^. In contrast, DNA-based approaches typically require substantially higher coverage to recover rare taxa. Importantly, the use of single-mosquito sequencing further preserved inter-individual microbiome heterogeneity that can be masked by pooled samples, improving the detection of rare and individual-specific microbial taxa and expanding the observed diversity of the mosquito-associated microbiome^13,31,46^. Together, these features likely underpin the high proportion of novel taxa identified here and demonstrate the power of meta-transcriptomics for revealing hidden microbial diversity.

Beyond overall diversity, our study provides a systematic and quantitative characterization of non-viral microbial communities at the level of individual hosts, enabling direct comparison of their abundance and co-occurrence. Bacterial communities were dominated by intracellular endosymbionts, particularly *Wolbachia* and Chlamydiota-related lineages. Although previous studies have shown that *Wolbachia* is widespread in mosquitoes, its detection status and relative abundance vary among mosquito species, often exhibiting pronounced host-associated differences^47–49^. Our single-mosquito meta-transcriptomic framework enabled direct comparisons of *Wolbachia* abundance and detection frequency across mosquito species, providing a more refined view of its ecological context in natural populations. The detection of Chlamydiota-like lineages further suggests the presence of previously unrecognized intracellular bacteria in mosquitoes^50,51^. In addition to these endosymbionts, we identified multiple taxa likely associated with the mosquito gut microbiota (e.g., *Serratia*^16^ and *Rosenbergiella*^40^), consistent with prior reports. Notably, several bacterial taxa identified here have been implicated in modulating arboviral infections. Although detected at low abundance, their presence suggests that functionally relevant microbes may be widespread, but underrepresented. Collectively, these findings indicate that bacterial communities are not only diverse and abundant, but also often host-specific and functionally relevant, highlighting their potential as targets for microbiome-based vector control strategies.

Fungi and other eukaryotic microbes, including Microsporidia and nematodes, constitute a diverse and ecologically heterogeneous component of the microbiome and may play important roles in shaping vector competence. Mechanistically, these organisms can influence pathogen susceptibility through both antagonistic and facilitative pathways^18,38,52^. Microsporidians may suppress infection via competition for host resources or immune priming^38,53^, and we observed that *Anopheles sinensis* harbors markedly lower microsporidian loads than other species, raising the possibility of increased permissiveness to infection. Conversely, facilitative effects have also been reported; for example, a gut-associated fungus (*Talaromyces* sp.) in *Aedes aegypti* enhances dengue virus infection by suppressing midgut trypsin activity, thereby promoting viral entry^18^. Hence, eukaryotic microbes may enhance pathogen invasion through modulation of host physiology. Nematodes, another abundant group, may similarly influence infection dynamics via effects on host metabolism or immune homeostasis, although their roles remain to be elucidated^54–56^. Together, these findings highlight eukaryotic microbes as underappreciated regulators of vector competence^57^.

From a public health perspective, our results are consistent with previous observations that arboviruses and other vector-borne pathogens represent only a small fraction of the overall mosquito microbiome^13,57^. This is expected, as pathogen prevalence in natural mosquito populations is typically low under non-outbreak conditions^13,58,59^. Notably, despite sampling known vector species such as *Culex tritaeniorhynchus*, we did not detect Japanese encephalitis virus (JEV), further supporting the notion that arboviruses occur at low abundance in natural settings. We also identified several putative novel pathogens; however, these taxa were detected at low abundance and prevalence, and their capacity for transmission to humans remains uncertain as their presence in mosquitoes does not necessarily indicate vector competence^60^. Nevertheless, these findings underscore the importance of continued surveillance of arthropod-borne pathogens^61^. It is likely that pathogen prevalence increases during outbreak conditions, although it may still remain substantially lower than that of insect-specific microbes, which are often stably maintained at higher abundance within mosquito populations^13^.

A key conceptual advance of this study is the demonstration that the mosquito microbiome constitutes a highly interconnected, cross-kingdom microbial network. Positive associations among taxa were more prevalent than negative ones; however, most taxon pairs showed no significant association. Within this largely sparse interaction structure, the observed positive associations suggest that coexistence and potential facilitative interactions may be common features of mosquito-associated communities^62,63^. Antagonistic interactions were also evident, with several candidate antiviral taxa—including *Wolbachia* and other bacteria—negatively associated with viral abundance^14,34,36,40^. However, these interactions are context– and species-dependent; for example, *Wolbachia* has been reported to suppress some viruses^14,34^ while facilitating others^64,65^. Because our findings are derived from natural mosquito populations, they likely reflect long-term ecological interactions and highlight the importance of ecologically grounded microbiome strategies for vector-borne disease control.

This study has several limitations. First, uneven sampling density may bias regional comparisons. Second, meta-transcriptomic data primarily capture transcriptionally active organisms and may underestimate DNA viruses or low-expression taxa. Third, many detected microorganisms remain incompletely characterized, limiting functional interpretation. Finally, inferred interaction networks are statistical and require experimental validation to establish causality.

In sum, this study provides a comprehensive, multi-kingdom framework of the mosquito microbiome, revealing its diversity, ecological structure, and interaction landscape. Beyond cataloging microbial diversity, our findings highlight mosquitoes as dynamic ecological systems shaped by interacting microorganisms and establish a large-scale baseline data set for future studies on vector competence, pathogen emergence, and microbiome-based disease control.

## Materials and Methods

### Sample collection

Between 2018 and 2025, more than 200,000 individual mosquitoes were collected across China, spanning 23 province-level regions, including Yunnan, Xinjiang, Tibet, Heilongjiang, and Inner Mongolia. Mosquitoes were captured using ultraviolet light traps deployed overnight (approximately 12 hours) at each sampling site, or by handheld collectors when appropriate. Following collection, mosquitoes were temporarily immobilized on dry ice and subjected to preliminary morphological identification by trained field biologists. Specimens were then sorted, transferred into cryogenic vials, and preserved on dry ice or in liquid nitrogen during transportation. All samples were subsequently stored in a −80°C freezer upon arrival at the laboratory until RNA extraction. From the full collection, a total of 5,163 individual mosquitoes were selected for downstream experiments and meta-transcriptomic analysis, based on sampling time, location, morphologically identified species, and habitat representation (Table S1). Of these, 2,426 individuals were included in a previous virome study using the same sequencing strategy, such that the present study incorporates an additional 2,737 individuals, substantially expanding geographic coverage across 15 provinces and including 55 mosquito species.

### Sample processing, RNA extraction, and sequencing

Prior to RNA extraction, each mosquito was washed three times with 1 mL of sterile, RNA– and DNA-free phosphate-buffered saline (PBS; Gibco, C10010500BT) to minimize external microbial contamination. Specimens were then individually homogenized in 600 µL of lysis buffer using a TissueRuptor (Qiagen, 9002756). Total RNA was extracted using the RNeasy Plus Mini Kit (Qiagen, 74104) following the manufacturer’s instructions, and RNA quality and integrity were assessed using an Agilent 2100 Bioanalyzer (Agilent Technologies). For each mosquito, total RNA was used to construct DNA nanoball (DNB)-based sequencing libraries with the MGIEasy RNA Library Prep Kit V3.0. Libraries were sequenced using paired-end 150 bp reads on the DNBSEQ T series platform (MGI, Shenzhen, China), generating individual-level meta-transcriptomic data.

### Mosquito species identification

Following preliminary morphological identification, species assignments were further confirmed using the cytochrome c oxidase subunit I (*COX1*) gene. From the assembled contigs of each library, candidate *COX1* sequences were first identified by DIAMOND v2.0.15^66^ blastx searches against reference *COX1* sequences downloaded from the NCBI RefSeq database (e-value ≤ 1 × 10⁻⁵). These candidates were subsequently queried against the NCBI non-redundant (nr) protein database using the same parameters, and contigs matching mosquito *COX1* sequences were retained. The retained contigs were trimmed to the *COX1* region to obtain high-confidence sequences. Non-rRNA reads were then mapped back to the curated *COX1* sequences to reduce potential assembly errors. The resulting sequences were submitted to the Barcode of Life Data System (BOLD)^67^ for species-level identification and further validated through phylogenetic analysis. Phylogenetic trees were estimated using the maximum likelihood (ML) method available in IQ-TREE2 v2.2.2.7^68^ with automatic model selection (ModelFinder Plus)^69^, and species assignments were confirmed based on tree topology.

Libraries showing evidence of mixed-host contamination were excluded from downstream analyses. Specifically, libraries were considered contaminated if multiple mosquito-species *COX1* sequences were detected in the same library, or if both mosquito-derived and non-mosquito insect-derived *COX1* sequences were identified.

### Virus identification

For each individual mosquito library, raw meta-transcriptomic reads were first processed to remove sequencing adapters and low-quality bases using fastp v0.20.1^70^. Duplicate reads and low-complexity reads were subsequently filtered using SOAPnuke v2.1.5^71^ and PRINSEQ++ v1.2^72^, respectively. Quality-controlled reads were then mapped to the SILVA rRNA database (release 138.1)^73^ using URMAP v1.0.1480^74^ and Bowtie2 v2.3.5.1^75^ to remove rRNA sequences. To eliminate host-derived contamination, non-rRNA reads were further mapped against mosquito reference genomes downloaded from NCBI, yielding non-host reads.

Clean reads were assembled *de novo* into contigs using MEGAHIT v1.2.8^76^ with default parameters and a minimum contig length of 300 bp. Assembled contigs were annotated against the NCBI nr database using DIAMOND blastx with an e-value threshold of 1 ×10^−5^. Taxonomic assignments were based on NCBI taxonomic identifiers (taxids) derived from blastx results, and virus-associated contigs were extracted for downstream analyses. Contigs associated with endogenous viral elements or other non-viral sources were removed.

Putative viral contigs were further validated based on the presence of conserved viral marker genes: the RNA-dependent RNA polymerase (RdRP) for RNA viruses and the non-structural protein 1 (NS1), DNA polymerase (DNApol), or replication-associated protein (Rep) for DNA viruses. Viral contigs shorter than 600 bp were excluded. For viruses with incomplete genomes, sequences were required to span the full conserved protein motif (e.g., the PALM domain of RdRP) to be considered valid viral representatives, and this was manually verified by multiple sequence alignment. Viral supergroups used in the downstream phylogenetic and diversity analyses refer to deep evolutionary lineages of RNA viruses defined based on RdRP-based phylogenetic relationships^7^.

### Definition of mosquito-associated viruses

Mosquito viromes comprise viruses associated with mosquitoes as well as those derived from other microbial taxa present within the host. To focus on viruses relevant to mosquito biology, we defined mosquito-associated viruses as those with the potential to infect mosquitoes or vertebrate hosts, distinguishing them from viruses likely associated with other organisms (e.g., fungi, nematodes, or bacteria). Accordingly, mosquito viruses were classified based on three criteria: sequence similarity to viruses with known arthropod– or vertebrate-associated hosts; high abundance within individual mosquitoes (≥1,000 RPM); phylogenetic placement within established arthropod-associated viral clades.

### Bacteria and archaea identification

To identify bacterial and archaeal community members, non-rRNA reads were first screened against marker gene references from the Genome Taxonomy Database (GTDB release 220)^77^. Specifically, non-rRNA reads were compared against bacterial *rpoB* (DNA-directed RNA polymerase β subunit; TIGR02013) and archaeal *rpoA1* (DNA-directed RNA polymerase subunit A′; TIGR02390) using DIAMOND blastx (e-value ≤ 1 × 10⁻²). Reads matching either marker gene were extracted and de novo assembled into contigs using MEGAHIT with default parameters and a minimum contig length of 300 bp. Contigs generated from marker-matching reads were combined with contigs assembled from clean reads of individual libraries before downstream marker-gene screening. Candidate marker-associated contigs were translated into protein sequences using ORFfinder v0.4.3^78^ with the appropriate genetic code (NCBI Translation Table 11). Candidate ORF protein sequences were subsequently deduplicated using CD-HIT v4.8.1^79^ and compared against GTDB *rpoB* and *rpoA1* reference sequences using DIAMOND blastp. To confirm marker gene identity, translated proteins were further screened using Hidden Markov Model (HMM)^80^ searches based on profiles constructed from aligned GTDB reference sequences. Only ORFs containing conserved *rpoB*– or *rpoA1*-associated domains were retained. These validated marker sequences were subsequently compared against the NCBI nr protein database using DIAMOND blastp, and final taxonomic assignments were determined based on sequence similarity and phylogenetic placement (see below).

### Fungi and other eukaryotic microbes identification

Identification of fungi and other eukaryotic microbes followed a pipeline analogous to that used for bacterial and archaeal discovery, with the translation elongation factor 1-alpha (*TEF1*) employed as a conserved and broadly expressed marker gene for eukaryotic taxa.

### Phylogenetic analyses

Phylogenetic analyses were performed to support taxonomic assignments at both the species and higher taxonomic levels (e.g., phylum and class). To infer the evolutionary relationships of the viruses identified, phylogenetic analyses were performed based on amino acid sequences of conserved viral hallmark proteins, comprising the RdRP for RNA viruses and NS1, DNApol, or Rep proteins for DNA viruses. For each viral supergroup, representative reference sequences spanning known diversity and those with the closest BLAST hits to the identified sequences were retrieved and included in the analyses.

For bacteria, archaea, fungi, and other eukaryotic microbes, phylogenetic analyses were based on amino acid sequences of appropriate marker genes: *rpoB* (bacteria), *rpoA1* (archaea), and *TEF1* (fungi and other eukaryotes). Reference sequences for *rpoB* and *rpoA1* were obtained from the Genome Taxonomy Database (GTDB), whereas *TEF1* references were retrieved from NCBI.

Amino acid sequences were aligned using MAFFT v7.505^81^ (E-INS-i algorithm), and ambiguously aligned regions were trimmed using TrimAl v1.4.rev22^82^. Alignment quality was assessed by visual inspection. Phylogenetic trees were estimated using the ML method in IQ-TREE2, with optimal substitution models selected by ModelFinder Plus (MFP). The resulting phylogenetic trees were then annotated and visualized using the ggtree package implemented in R.

### Species demarcation criteria

Species-level delineation was performed based on phylogenetic placement and amino acid sequence identity thresholds derived from conserved marker genes for each microbial group. For viruses, classification was based on the RdRP for RNA viruses and NS1, DNApol, or Rep for DNA viruses. Viral lineages were assigned to known species if they shared ≥90% amino acid identity together with either ≥80% nucleotide identity or ≥80% alignment coverage, or if they had <90% amino acid identity but ≥90% nucleotide identity with ≥80% alignment coverage; all others were considered putatively novel viral species.

For cellular microbes, species delineation was based on marker genes specific to each taxonomic group. Bacterial taxa were defined using *rpoB* (DNA-directed RNA polymerase β subunit), and archaeal taxa using *rpoA1* (RNA polymerase subunit A′), both with a threshold of 98% amino acid identity. Fungi and other eukaryotic microbes were delineated based on translation elongation factor 1-alpha (*TEF1*) sequences, using a more stringent threshold of 99% amino acid identity to account for higher conservation within eukaryotic lineages.

### Full-spectrum microbiome quantification: coarse compositional profiling across microbial categories

Here, the mosquito full-spectrum microbiome refers to the transcriptome-detectable ensemble of viruses, bacteria, archaea, fungi, and other eukaryotic microbes associated with mosquitoes. To capture the overall composition of the mosquito full-spectrum microbiome, we first performed a coarse, global-level quantification, in which all assembled contigs and unassembled reads were assigned to taxonomic categories spanning the host (mosquito), RNA and DNA viruses, bacteria, archaea, fungi, and other eukaryotic microbes. Assembled contigs were annotated against the nr and nt databases, followed by taxonomic annotation of the remaining unassembled reads. To refine abundance estimation, non-rRNA reads were subsequently mapped back to the annotated contigs, and read counts were aggregated from both contig-based read mapping and direct read-level annotations. Integration of these two sources generated a composite taxonomic profile, providing a comprehensive and quantitative overview of full-spectrum microbiome composition across all microbial groups.

### Full-spectrum microbiome quantification: species-level resolution

To obtain higher-resolution, species-level estimates of full-spectrum microbiome composition, we next performed a targeted quantification based on reference mapping. Prior to quantification, a standardized quality control and filtering pipeline was applied to all sequences detected. Host-derived sequences were first removed to minimize contamination. To assess potential assembly errors, non-host reads were mapped back to assembled viral genomes or microbial marker genes. Species-level quantification was then performed using reference-based read mapping. Non-host reads were aligned to target sequences using Bowtie2, with different strategies applied to distinct microbial groups. RNA viruses were quantified by mapping reads to complete or near-complete assembled viral genomes, whereas DNA viruses were quantified using either full genomes or core transcript regions depending on assembly completeness, with viral genomes or marker genes covered by <300 bp excluded. In contrast, bacteria, archaea, fungi, and other eukaryotic microbes were quantified based on marker gene–derived sequences, reflecting the limited recovery of complete genomes for most cellular taxa. Abundance-based filtering was then applied to reduce false positives: taxa were excluded if their read counts were <0.1% of the maximum observed for the same taxon within the same sequencing lane (i.e., to mitigate index hopping), or if their abundance was extremely low (RPM < 1). Abundance was calculated as the number of reads per million non-rRNA reads (RPM) for each taxon and used for downstream analyses.

### Vertebrate pathogen identification

Putative vector-borne pathogens that are likely to infect humans and other mammals were identified based on a combination of phylogenetic placement and sequence similarity criteria. For vector-borne viruses, classification followed the phylogenetic framework described above. Viral sequences sharing ≥90% amino acid identity with known vector-borne viruses, or clustering within established vector-borne viral clades, were classified as putative vector-borne viruses.

For bacterial pathogens, phylogenetic analyses and sequence comparisons were conducted based on *rpoB* nucleotide sequences. Bacterial taxa sharing ≥97% nucleotide identity with reference pathogens listed in the National Catalogue of Human Infectious Pathogens (https://www.nhc.gov.cn/qjjys/c100016/202308/57f94fa7e121475ab87ff0ea14a92f6c.shtml) were classified as known bacterial pathogens, a criterion used in *rpoB* amplicon-based OTU clustering^83^, whereas those exhibiting 95–97% nucleotide identity were designated as putative novel bacterial pathogens.

For fungal and parasitic pathogens, identification was based on *TEF1* protein sequence similarity together with phylogenetic analysis. Sequences assigned to the same species as known pathogenic taxa listed in the National Catalogue of Human Infectious Pathogens were classified as recognized fungal pathogens, whereas those sharing 95–99% identity with known pathogens were considered putative novel fungal pathogens. Parasite-associated sequences sharing ≥99% amino acid identity with reported species were classified as known parasitic pathogens, whereas those sharing 95–99% identity were classified as putative novel parasitic pathogens.

### Statistical analysis

All statistical analyses were conducted in R v4.4.1 using RStudio. To investigate potential interactions within the mosquito full-spectrum microbiome, Spearman’s correlation analysis was applied at the species level. To minimize confounding effects from non-mosquito hosts, only viruses classified as mosquito-associated (as defined above) were included.

For correlation analysis, associations were evaluated based on microbial abundance (RPM) across viruses, bacteria, archaea, fungi, and other eukaryotic microbes. Correlations were calculated only for taxon pairs co-detected in at least 10 libraries (i.e., both taxa had non-zero RPM values in ≥10 libraries), and pairs not meeting this criterion were excluded. Unless otherwise specified, associations with *p* < 0.05 were considered statistically significant and were retained as candidate microbial associations.

Phylogenetic trees were annotated and visualized using ggtree. Chord diagrams were generated using circlize (chordDiagram function), Venn plots were produced using ggvenn, and additional figures were generated using pheatmap and ggplot2.

## Acknowledgments

This work was supported by the National Key R&D Program of China (2024YFC2607504), the National Natural Science Foundation of China (32270160), the National Natural Science Foundation of China (81290342), the Prevention and Control of Emerging and Major Infectious Diseases-National Science and Technology Major Project (2025ZD01901104), the National Natural Science Foundation of China (32501494), the Fundamental Research Funds for the Central Universities, Sun Yat-sen University (24xkjc024), and the Major Project of Guangzhou National Laboratory (GZNL2023A01001). We thank the field teams, local Centers for Disease Control and Prevention, and collaborating institutions across China for assistance with mosquito collection, species identification, and sample transportation. We also thank the sequencing, data management, and computational support teams at Sun Yat-sen University, BGI Research, and the China National GeneBank for technical assistance.

## Author Contributions

Conceptualization, W.-C.W. and M.S.; methodology, Q.-Y.G., W.-C.W., P.-B.S., Z.-Q.D., D.-X.W., and M.S.; investigation, Q.-Y.G., W.-C.W., P.-B.S., G.-Y.L., Y.-F.P., J.W., Y.G., K.-J.L., H.-L.Z., S.-J.L., G.-Y.X., M.-W.P., Y.-Q.L., J.-M.H., C.-H.Y., S.-Q.M., J.-X.C., X.H., J.-B.K., X.-X.C., Z.-R.R., J.-H.L., and X.J.; resources (sampling), Q.-Y.G., W.-C.W., G.-Y.L., J.W., G.-Y.X., M.-W.P., Y.-Q.L., S.-Q.M., X.H., J.-B.K., X.-X.C., Y.F., K.L., W.-H.Y., D.W., B.Z., J.W., T.-Q.A., X.-Y.H., G.-D.L., and M.S.; formal analysis, Q.-Y.G., W.-C.W., and P.-B.S.; writing – original draft, Q.-Y.G. and M.S.; writing – review and editing, all authors; supervision, Z.-Q.D., M.S., and D.-X.W.; funding acquisition, Z.-Q.D., M.S., and D.-X.W.

## Ethics Statement

No human participants, vertebrate animals, or endangered or protected species were involved. All procedures complied with national and institutional regulations governing non-vertebrate field sampling, and formal ethics approval was not required.

## Conflict of Interest

The authors declare no competing interests.

## Data and Code Availability

● Raw meta-transcriptomic sequencing data have been deposited in the CNSA (CNGB Sequence Archive) of CNGBdb (China National GeneBank database), project accession: CNP0005544.
● Assembled viral contigs and taxonomic annotations are available in the China National GeneBank database (CNGBdb) under the accession N_AAKCIK010000000– N_AAKDPR010000000.
● Microbial marker-gene sequences and taxonomic annotations are available at figshare.
● Custom R scripts used for data analysis and visualization are available at figshare.
● Any additional information required to reanalyze the data reported in this paper is available from the corresponding author upon reasonable request.

## Notes

### Competing Interest Statement

The authors have declared no competing interest.

