## Supplemental figures for "Multi-kingdom microbial diversity and interaction landscapes in mosquitoes revealed by 5,163 individual meta-transcriptomes"

**List of Supplemental Materials**

- Figure S1. Phylogenetic relationships and sample sizes of mosquito species
- Figure S2. Phylogenetic relationships of the mosquito viromes
- Figure S3. Phylogenetic relationships of the bacteria sampled
- Figure S4. Characterization of the diversity and composition of prokaryotic communities
- Figure S5. Phylogenetic relationships of the eukaryotes sampled
- Figure S6. Comparative analysis of eukaryotic community diversity and prevalence among dominant mosquito species
- Figure S7. ML phylogenetic trees of pathogens identified in mosquitoes
- Figure S8. Interactions associated with Wolbachia

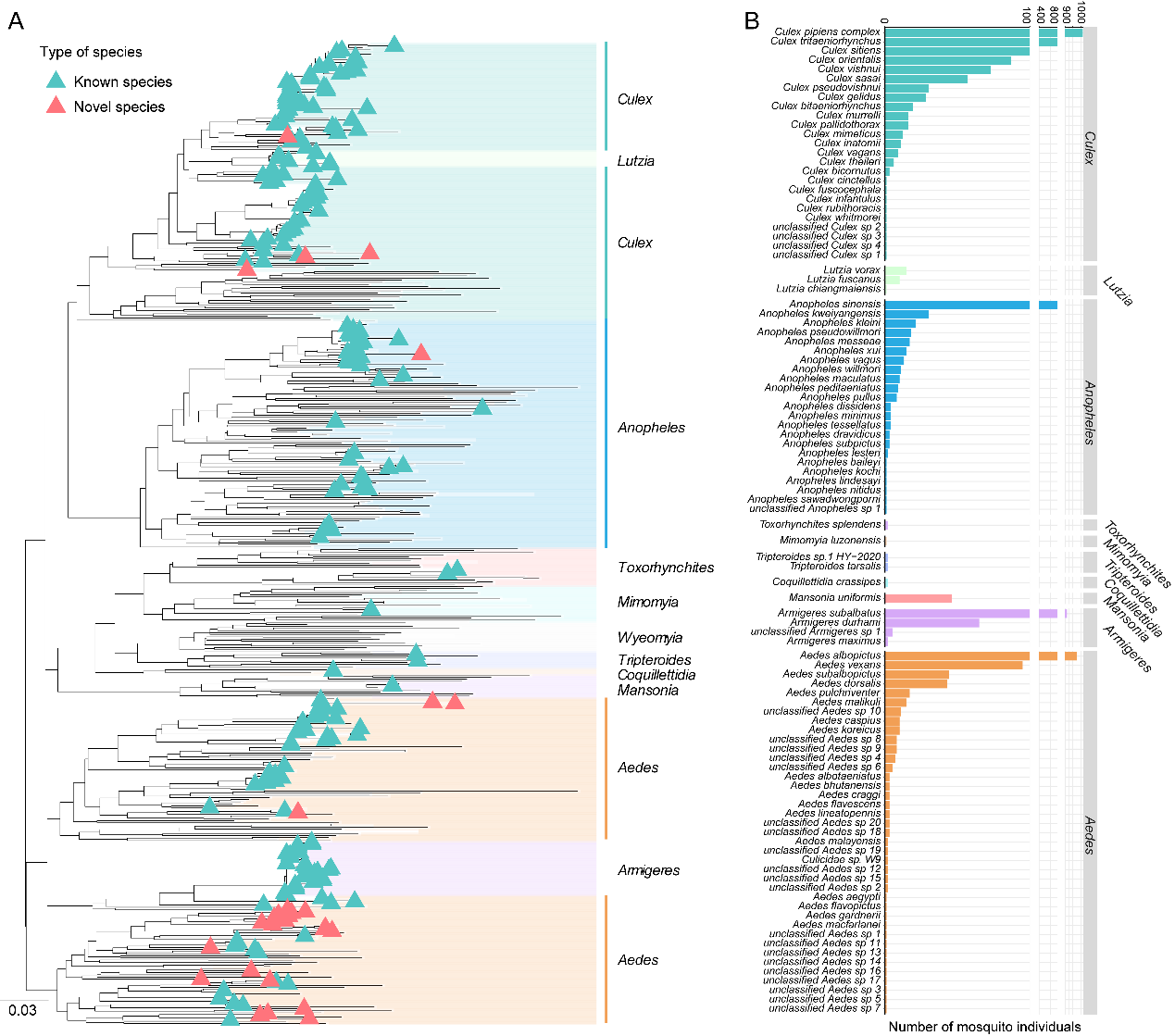

**Figure S1. Phylogenetic relationships and sample sizes of mosquito species.**

**(A)** Maximum likelihood phylogenetic tree of 100 mosquito species based on the mitochondrial *COX1* gene. Colored annotation bars indicate mosquito genera, which are also labeled on the right. Triangles indicate the representative *COX1* sequences of mosquito species identified in this study, with cyan denoting known species and red denoting novel species. **(B)** Sample size of each mosquito species, arranged in descending order by number of samples. Colors represent different mosquito genera.

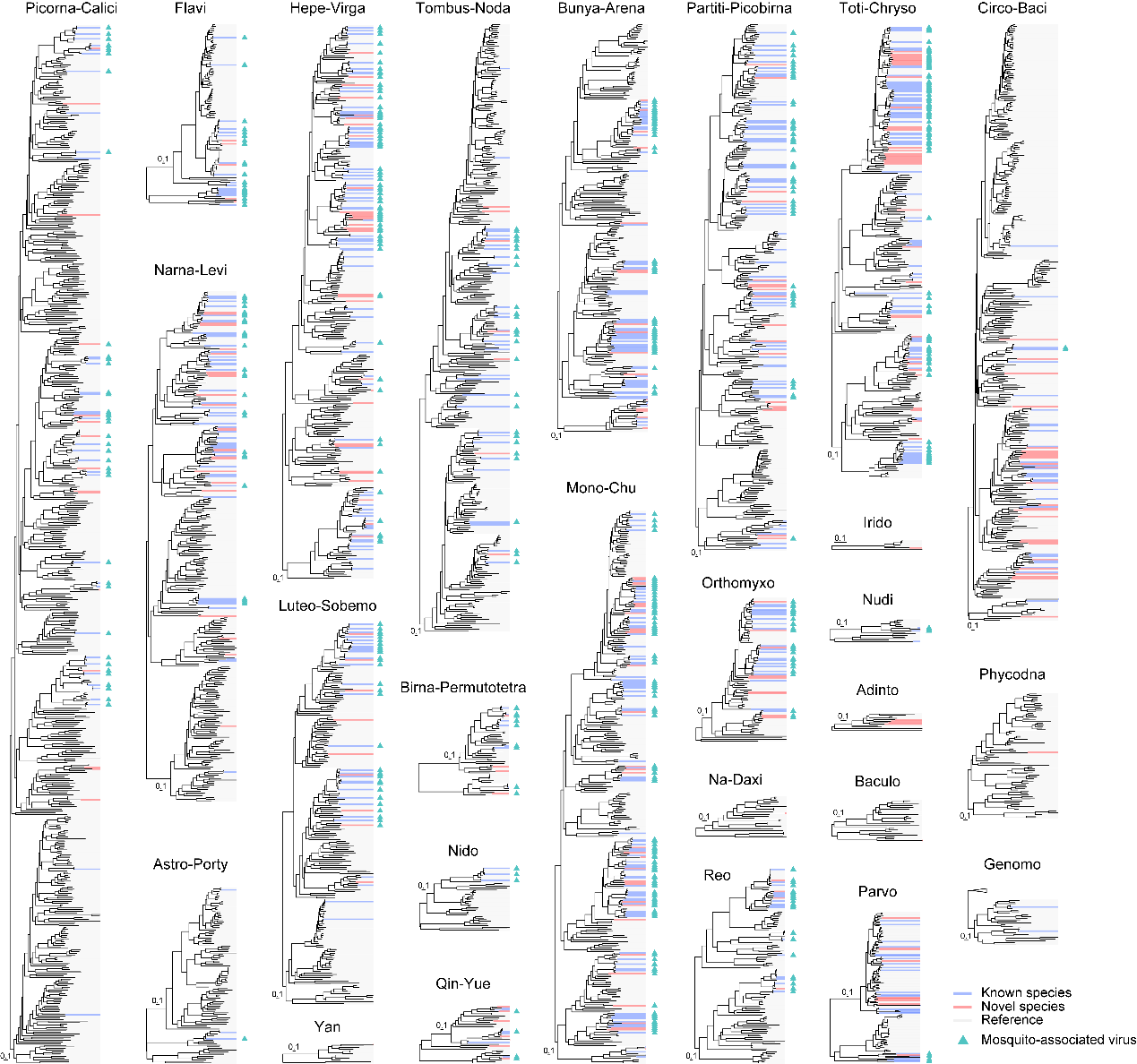

**Figure S2. Phylogenetic relationships of the mosquito viromes.**

ML phylogenetic trees of (i) RNA viruses estimated using the RNA-dependent RNA polymerase (RdRP) gene, and (ii) DNA viruses inferred using DNA polymerase, replication-associated protein (Rep), or non-structural protein 1 (NS1) genes. Known viral species are shown in blue, newly identified viruses in red, and mosquito-associated viruses are marked with triangles. Reference sequences are shown in gray.

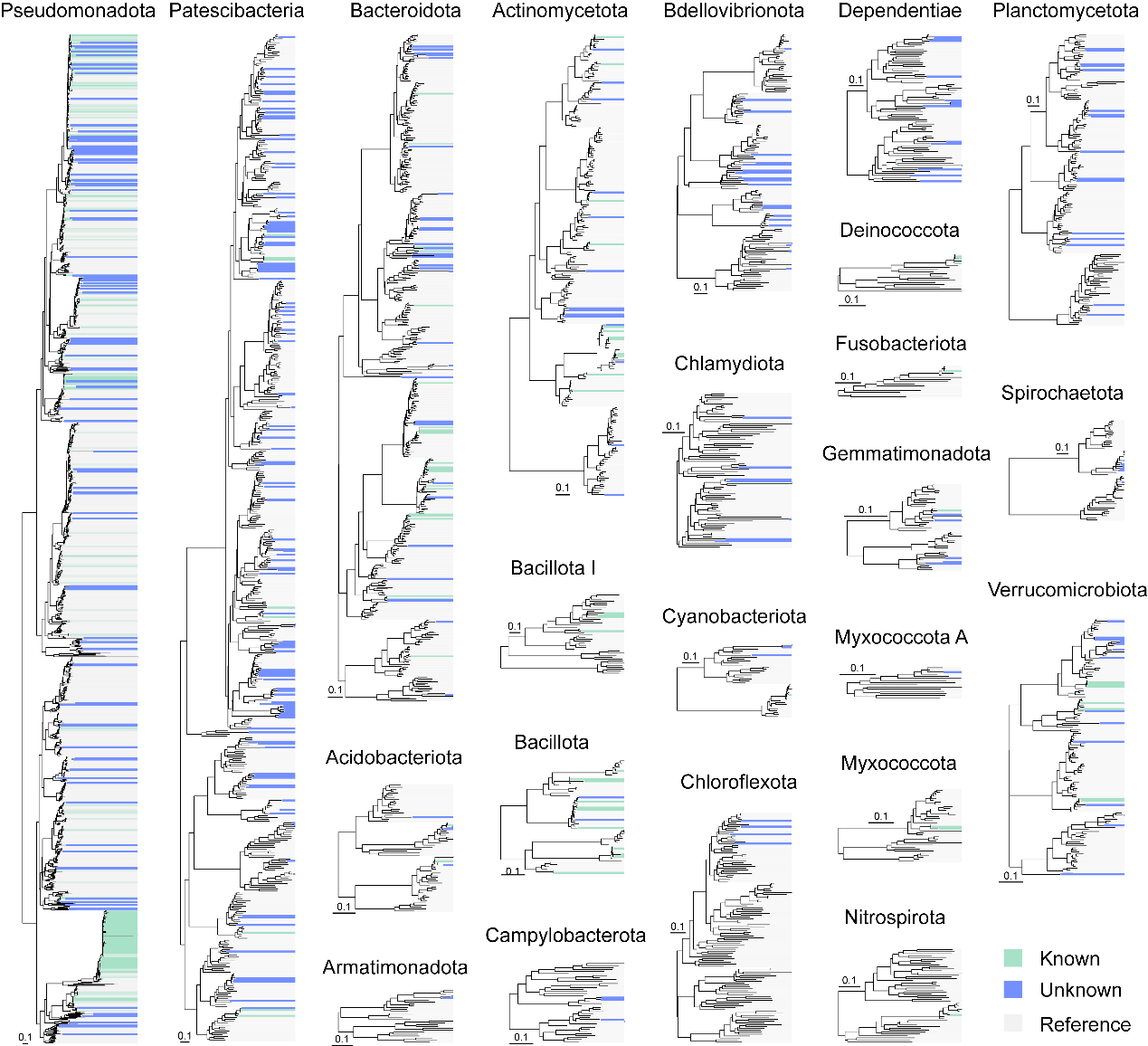

**Figure S3. Phylogenetic relationships of the bacteria sampled.**

ML phylogenetic trees of bacteria at the phylum level were inferred based on *rpoB* protein sequences. Known bacterial taxa are shown in green, putatively novel bacterial taxa are shown in blue, and reference sequences are shown in gray.

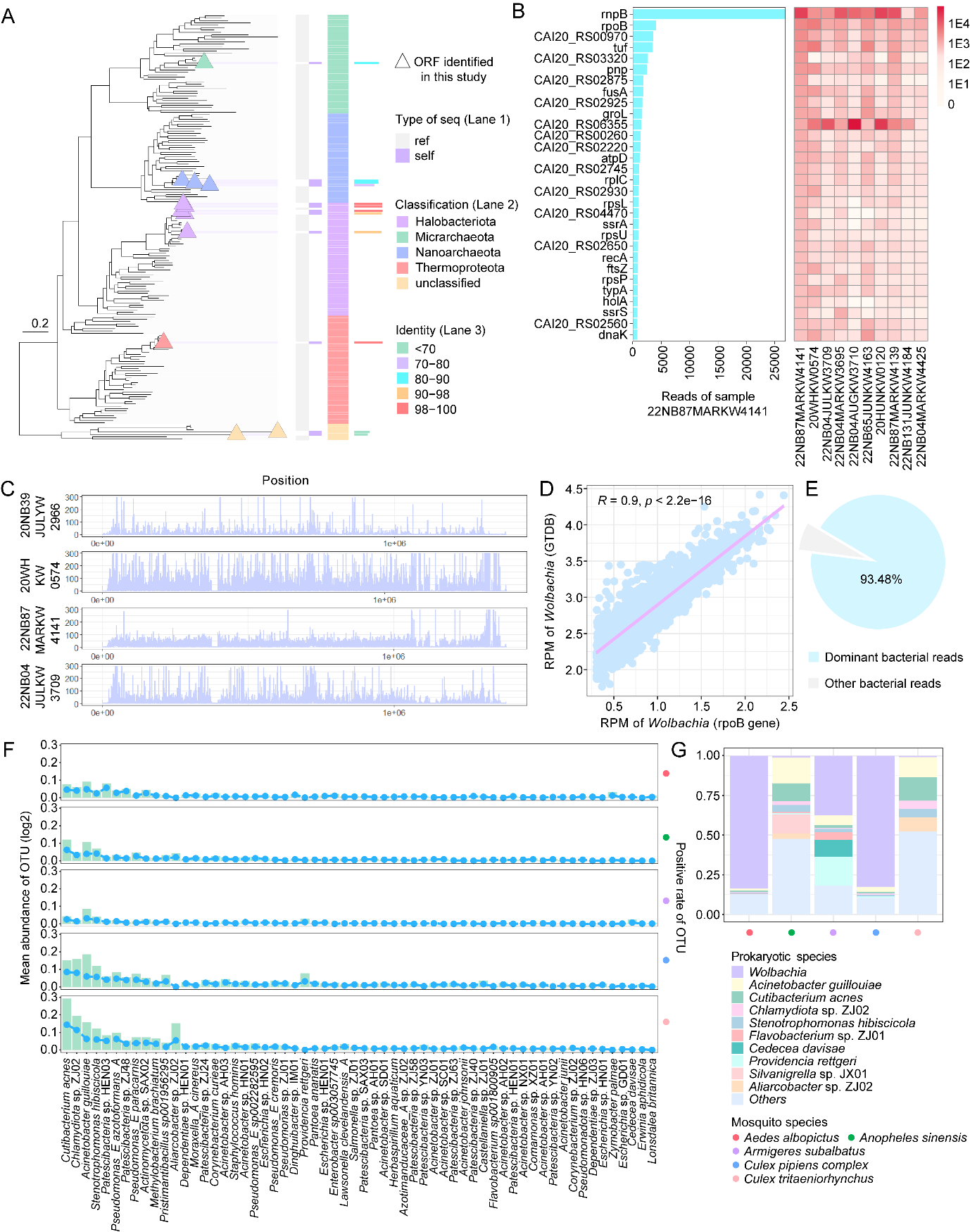

**Figure S4. Characterization of the diversity and composition of prokaryotic communities.**

**(A)** Archaeal phylogeny inferred using the *rpoA1* protein. The ML tree shows archaeal *rpoA1* sequences identified in this study (triangles), colored by phylum. From the inner to the outer rings, annotation tracks indicate sequence origin (reference vs. sequences identified in this study), phylum-level taxonomic assignment, and amino acid sequence identity, with each ring represented using a distinct color scale. **(B)** Gene-level abundance profiles of *Wolbachia* across individual samples. Heatmap showing the abundance of the most abundant *Wolbachia* genes across 10 representative samples. Rows correspond to individual genes and columns to samples, with color indicating abundance level. The bar plot on the left shows the gene-level abundance in one representative sample. **(C)** Genome-wide sequencing depth of coverage across the *Wolbachia* genome. **(D)** Correlation between *Wolbachia* abundance estimated using *rpoB*-based marker-gene quantification and genome-wide coverage-based methods. **(E)** Relative contribution of dominant bacterial taxa to total bacterial reads across all samples. **(F)** Abundance (left axis, bars) and overall prevalence (right axis, dots and dashed line) of dominant bacterial taxa across mosquito species. **(G)** Relative abundance of prokaryotic species across the five most abundant mosquito species. Colors denote prokaryotic species, and colored circles at the bottom denote mosquito species.

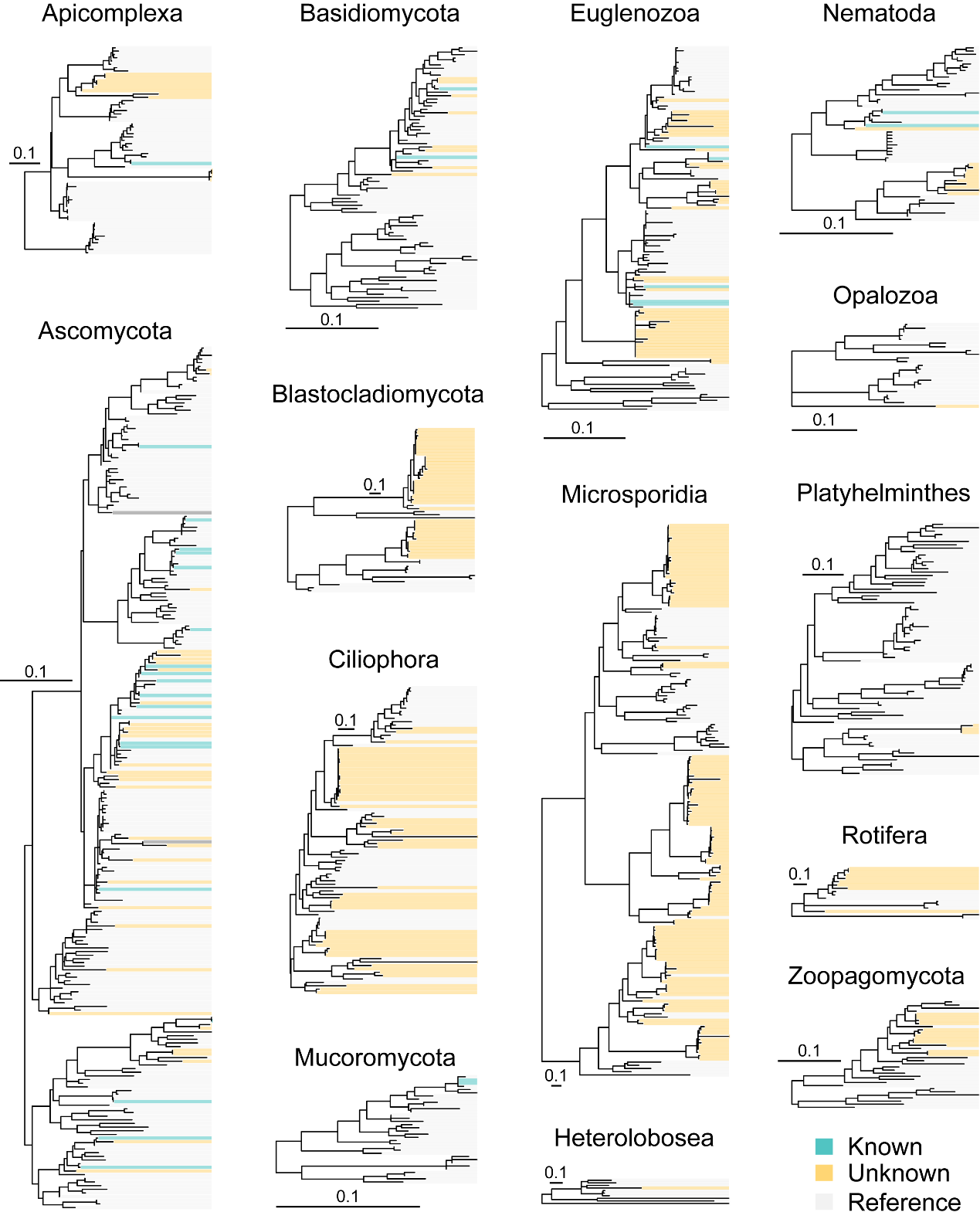

**Figure S5. Phylogenetic relationships of the eukaryotes sampled.**

ML phylogenetic trees of eukaryotic taxa at the phylum level were inferred from *TEF1* protein sequences. Known eukaryotic taxa are shown in blue, putatively novel taxa identified in this study are shown in yellow, and reference sequences are shown in gray.

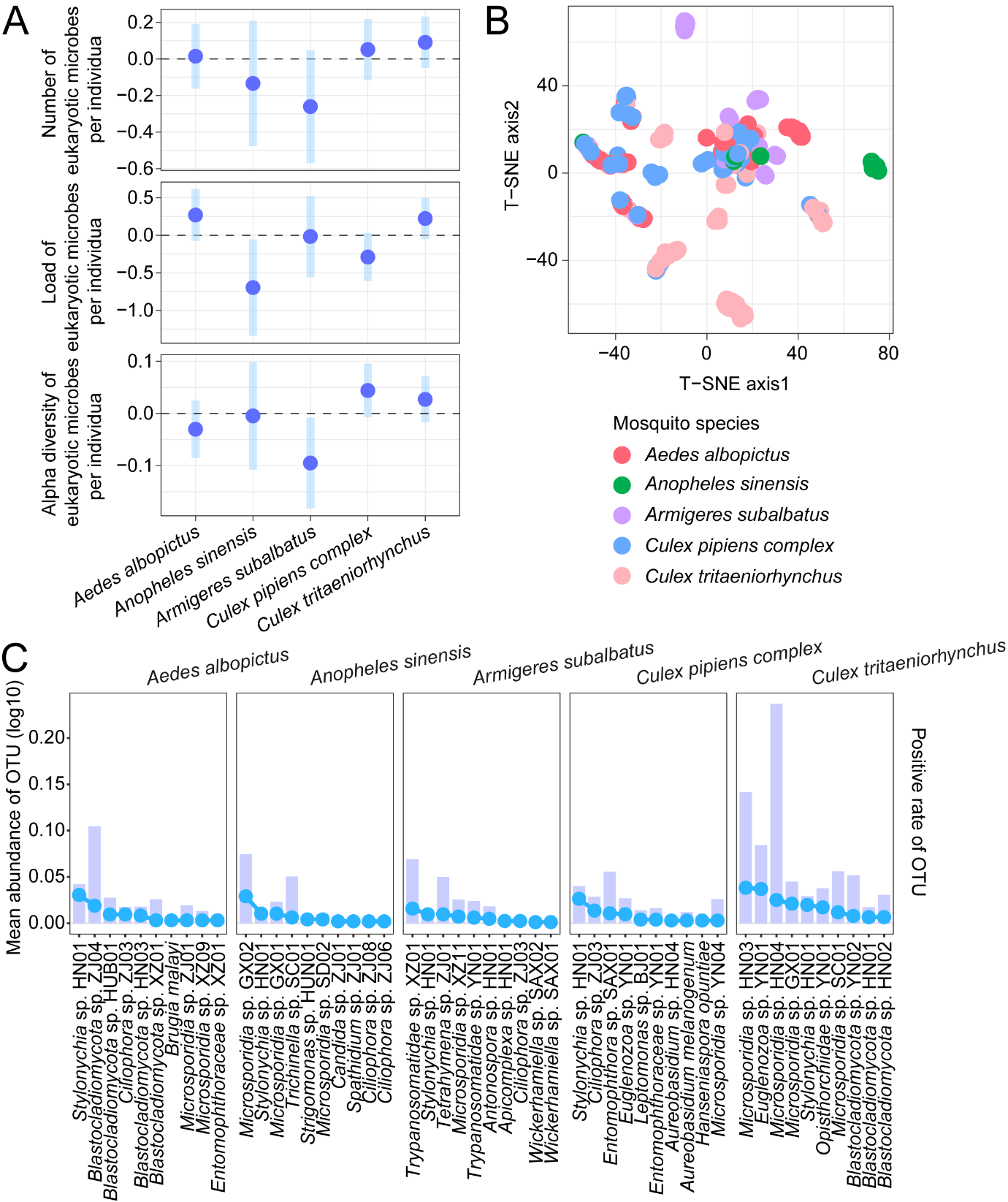

**Figure S6. Comparative analysis of eukaryotic community diversity and prevalence among dominant mosquito species.**

**(A)** Comparisons of eukaryotic microbial richness, load, and alpha diversity among the five dominant mosquito species. Points indicate model-estimated species effects, and bars indicate approximate 95% confidence intervals. **(B)** t-SNE ordination of individual mosquito eukaryotic microbial taxa composition. Each point represents one mosquito sample, colored according to mosquito species. **(C)** Mean abundance (left axis, bars) and overall prevalence (right axis, dots and dashed line) of the dominant eukaryotic taxa across dominant mosquito species.

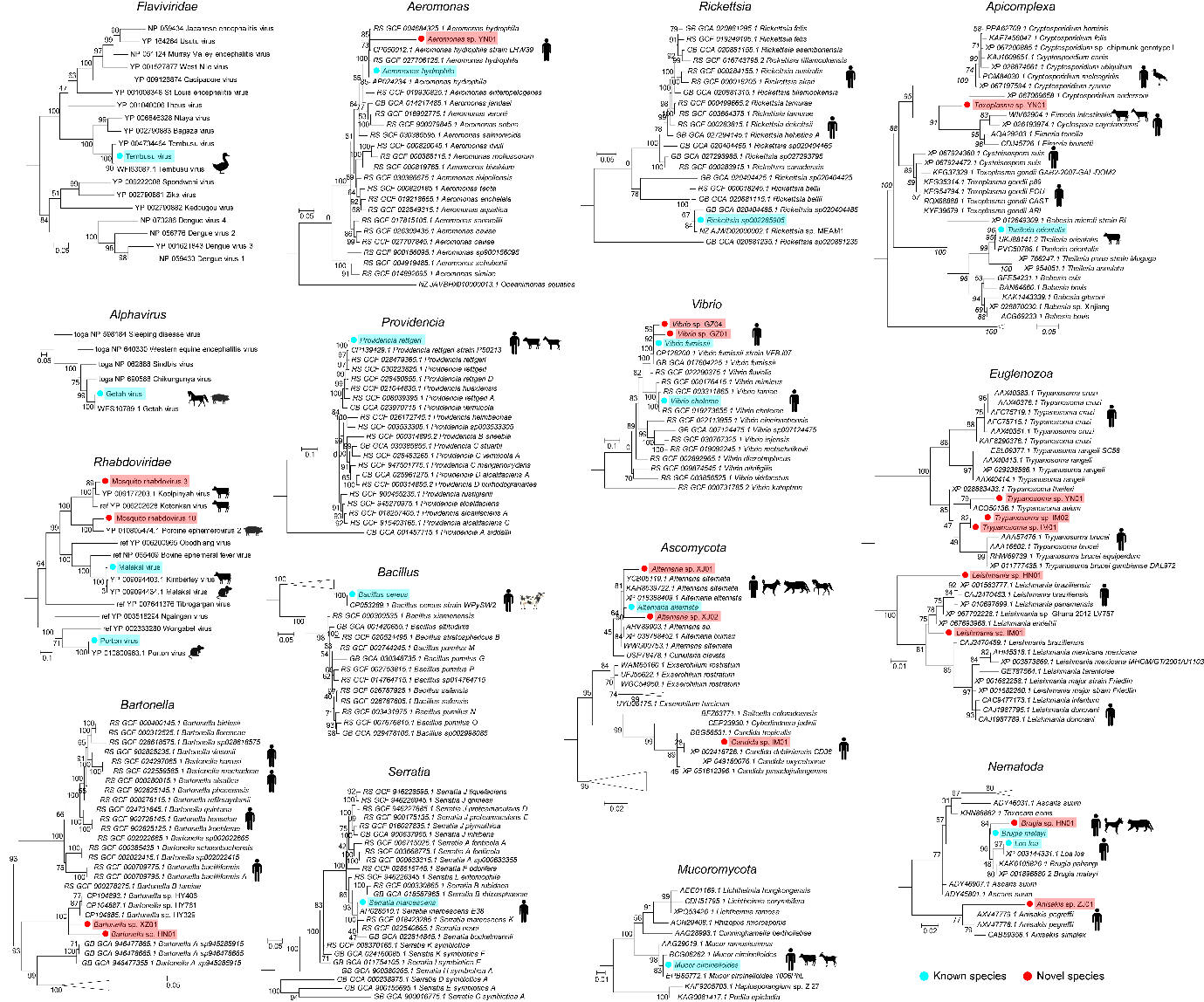

**Figure S7. ML phylogenetic trees of pathogens identified in mosquitoes.**

Amino acid sequence phylogenetic trees of arboviruses were inferred from the RdRP gene, bacterial phylogenies were estimated using the *rpoB* nucleotide sequences, and fungal and parasitic phylogenies were estimated from the *TEF1* protein sequences. Cyan circles and shaded areas indicate known pathogen-related taxa, whereas red circles and shaded areas indicate putative novel pathogen-related taxa identified in this study.

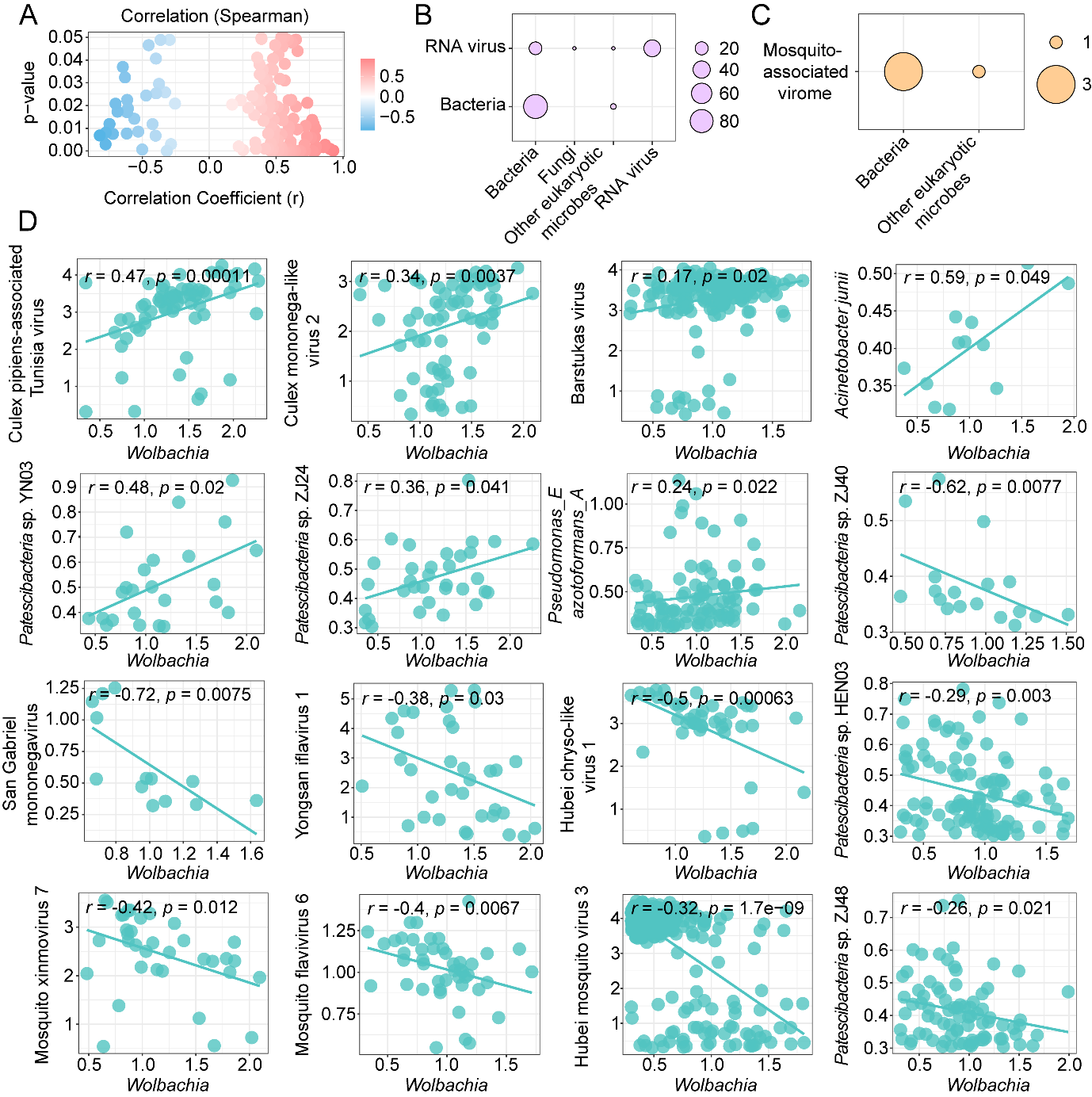

**Figure S8. Interactions associated with *Wolbachia*.**

**(A)** Distribution of Spearman correlation coefficients. Point colors indicate Spearman correlation coefficients (corresponding to the x-axis). **(B)** Number of interactions among different microbial categories based on Spearman correlation. Bubble size represents the number of significant associations. **(C)** Number of microbial taxa significantly associated with the mosquito-associated virome based on Spearman correlation. Bubble size represents the number of significant associations. **(D)** Correlations between *Wolbachia* and other microbes (Spearman).
